# Regulation of protein abundance in genetically diverse mouse populations

**DOI:** 10.1101/2020.09.18.296657

**Authors:** Gregory R. Keele, Tian Zhang, Duy T. Pham, Matthew Vincent, Timothy A. Bell, Pablo Hock, Ginger D. Shaw, Steven C. Munger, Fernando Pardo-Manuel de Villena, Martin T. Ferris, Steven P. Gygi, Gary A. Churchill

## Abstract

Proteins constitute much of the structure and functional machinery of cells, forming signaling networks, metabolic pathways, and large multi-component complexes. Protein abundance is regulated at multiple levels spanning transcription, translation, recycling, and degradation to maintain proper balance and optimal function. To better understand how protein abundances are maintained across varying genetic backgrounds, we analyzed liver proteomes of three genetically diverse mouse populations. We observe strong concordance of genetic and sex effects across populations. Differences between the populations arise from the contributions of additive, dominance, and epistatic components of heritable variation. We find that the influence of genetic variation on proteins that form complexes relates to their co-abundance. We identify effects on protein abundance from mutations that arose and became fixed during breeding and can lead to unique regulatory responses and disease states. Genetically diverse mouse populations provide powerful tools for understanding proteome regulation and its relationship to whole-organism phenotypes.

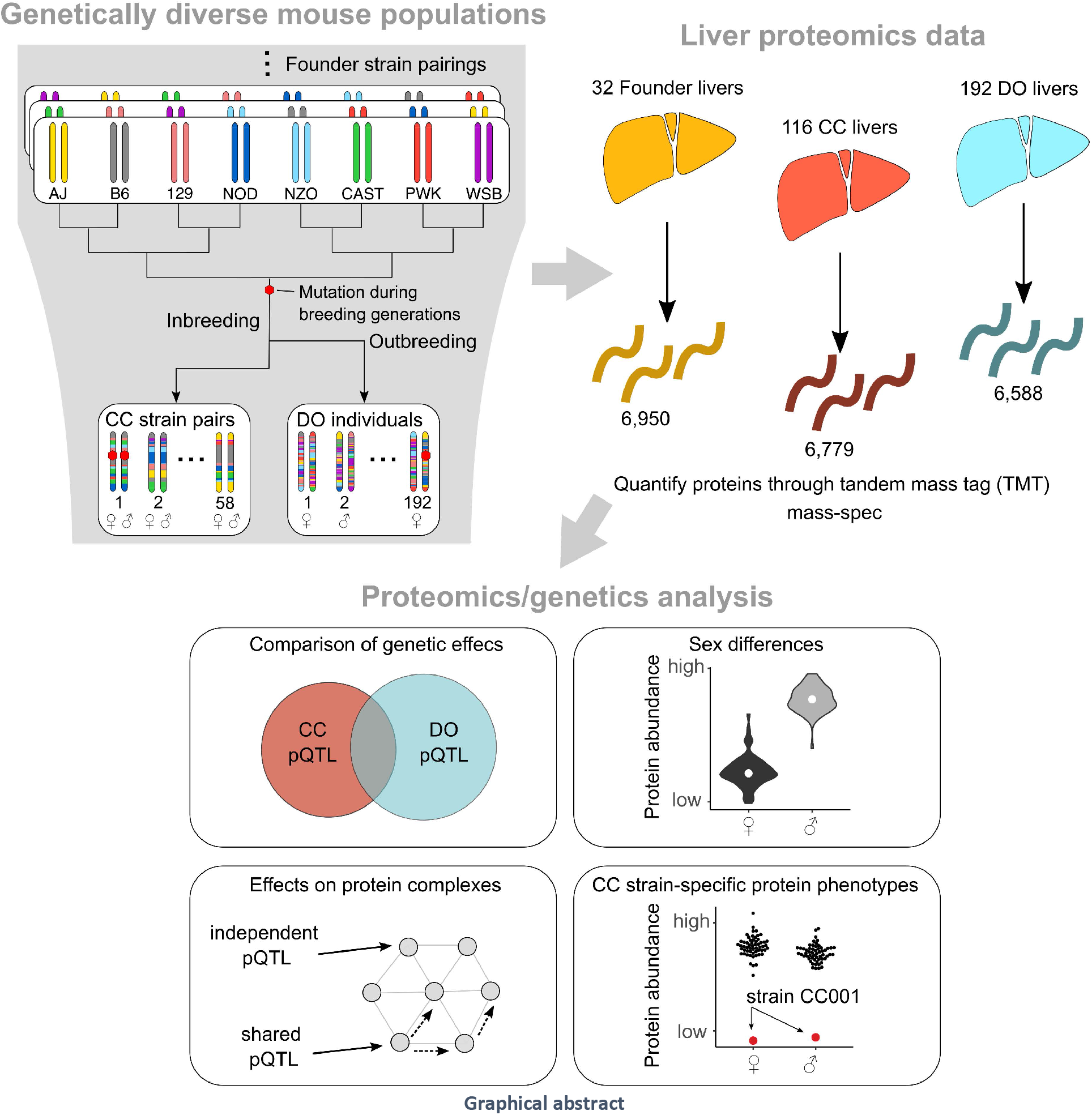

## INTRODUCTION

Protein abundance is regulated at multiple levels spanning transcription, translation, recycling, and degradation. It is responsive to genetic variation, as observed in yeast (Picotti et al., 2013), mice (Chick et al., 2016; Williams et al., 2016; Wu et al., 2014), and human cell lines, tissues, and populations (Battle et al., 2015; He et al., 2020; Liu et al., 2015; Suhre et al., 2020). Genetic effects on protein abundance can be broadly divided into two classes. Local variants occur in the vicinity of the coding gene and commonly affect protein abundance by altering the rate of transcription or stability of the transcript (Pai et al., 2012). In contrast, distal genetic variants are found at loci far from the coding gene and their effects on protein abundance are often post-translational and conferred via a diffusible intermediate; often another protein. Within these categories, multiple modes of regulation are possible (Chick et al., 2016). The complexity of genetic regulation is compounded for proteins that are members of complexes where stoichiometry imposes varying degrees of constraint (Chick et al., 2016; Huttlin et al., 2020; Romanov et al., 2019; Szklarczyk et al., 2019; Taggart et al., 2020). The number of genetic loci that affect a protein can range from a single locus (monogenic) to many (multi-genic to polygenic). Their individual effects can be additive or dominant/recessive, thus dependent on the extent of heterozygosity in the population. Loci can also interact, resulting in epistatic effects.

Resource populations with high levels of genetic diversity can help to identify and characterize the genetic loci that affect protein abundance. The Collaborative Cross (CC) (Churchill et al., 2004; Collaborative Cross Consortium, 2012) and Diversity Outbred (DO) (Churchill et al., 2012) mouse populations are powerful tools for genetic analysis. They are descended from a common set of eight inbred strains (*i.e.*, the founder strains), representing three subspecies of the house mouse, *Mus musculus* (Yang et al., 2007, 2011) and encompass genetic variation from across laboratory and wild mice. DO mice are available in large numbers and each individual possesses a unique, highly recombinant and heterozygous genome, supporting powerful, fine resolution mapping of genetic variants (*e.g.*, Svenson *et al.* 2012; French *et al.* 2015; Keller *et al.* 2018, 2019). The CC consists of ~60-70 strains that are largely inbred, with many strains homozygous at most loci (>99%), and residual heterozygous regions known and characterized (Collaborative Cross Consortium, 2012; Shorter et al., 2019; Srivastava et al., 2017). Due to fewer outbreeding generations than in the DO, the CC possess larger linkage disequilibrium (LD) blocks. The reproducible genomes of CC strains enable replicate study designs (Mosedale *et al.* 2017, 2019) and the characterization of genetic effects on strain-specific phenotypes (Philip *et al.* 2011; McMullan *et al.* 2016). CC strains can serve as models for human diseases including colitis (Rogala et al., 2014), susceptibility to Ebola infection (Rasmussen et al., 2014), influenza A virus (Noll et al., 2020), SARS-coronavirus (Gralinski et al., 2015), and peanut allergy (Orgel et al., 2019).

In this work, we obtained multiplexed mass spectrometry (mass-spec) quantification of proteins in liver samples of 116 CC mice representing female/male pairs from 58 strains. We previously collected proteomics data from the livers of 192 DO mice and 32 mice representing the eight founder strains (two animals of each sex per founder strain) (Chick et al., 2016) (**Figure 1a**). We map protein quantitative trait loci (pQTL) in both the CC and DO and find consistency of the genetic and sex effects on protein abundance across the populations. We characterize the genetic architecture of protein-complexes in both the CC and DO, and find that complexes with highly co-abundant members are heritable and exhibit non-additive and/or highly polygenic genetic architecture, which is more apparent in the CC. Finally, we identify genetic variants that originated and became fixed during the of the CC and contribute to CC strain-specific phenotypes. Our work demonstrates the complementary strengths of these genetic resources and offers new insights into the genetic regulation of protein abundance.

**Figure 1.**
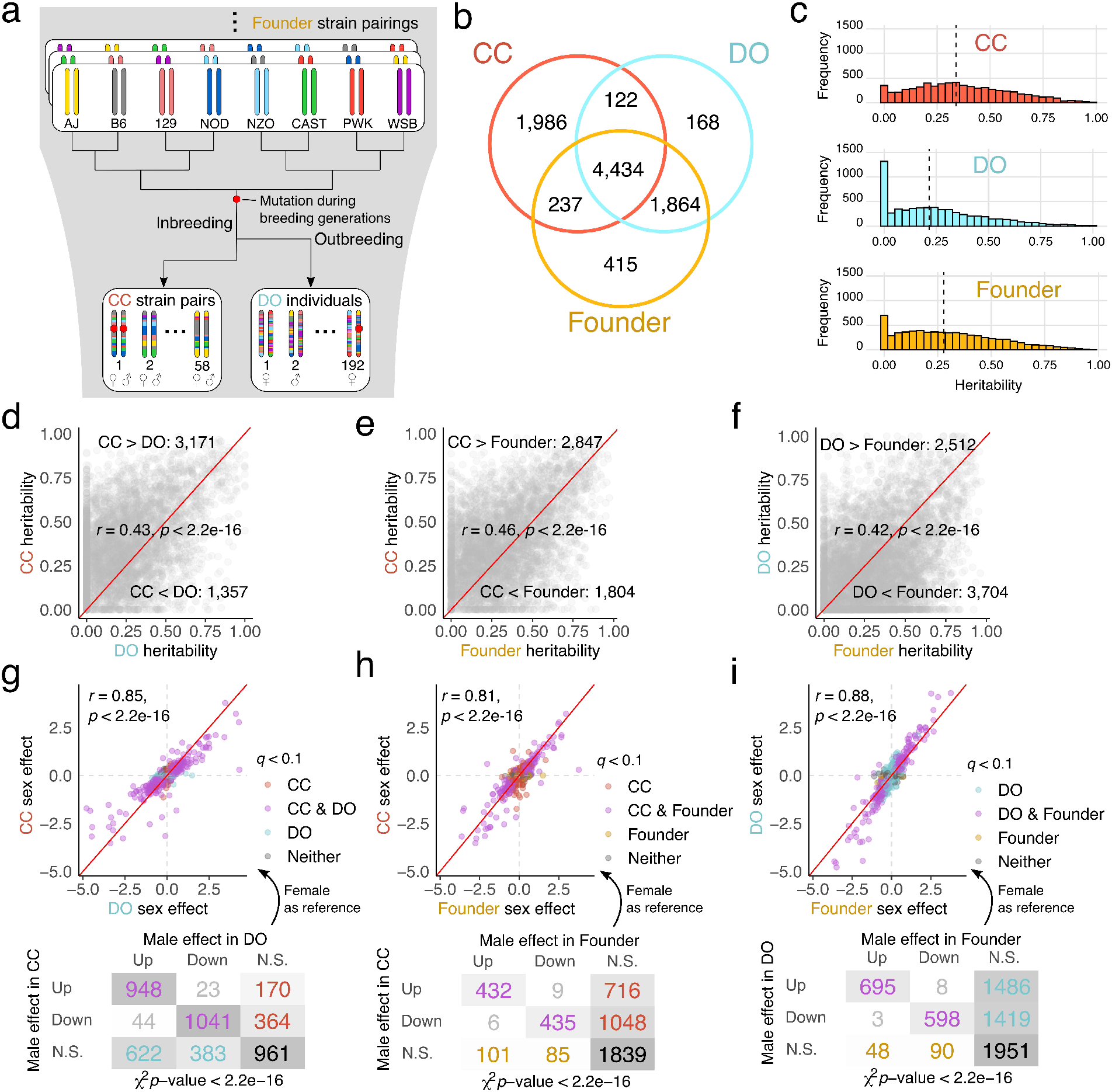
Comparisons of genetic and sex effects on protein abundance among the CC, DO, and founder strains reveal strong concordance. (a) The CC strains and DO mice are descended from the same eight inbred founder strains. Mutations occur during the breeding generations of the CC and DO and can become fixed in the CC, allowing their effects to be characterized. (b) Venn diagram of the analyzed proteins in the CC, DO, and founder strains. The founder strains and DO samples were from a joint experiment with the same protein annotations, resulting in greater overlap. (c) Overall genetic regulation, measured as heritability, is greater in the inbred CC and founder strains, reflecting contributions from non-additive genetic effects. Dashed vertical lines represent the median heritability in each population. Comparisons of the heritability of individual protein abundance for (d) CC and DO, (e) CC and founder strains, and (f) DO and founder strains. The correlation in heritability is high and significant regardless of which populations are being compared (*r* > 0.42, *p* < 2.2e-16). Heritability estimates in the CC are higher in part due to inclusion of a bridge sample across the batches of mass-spec experiments, which improved quantification accuracy (**Methods**). Red diagonal lines included for reference. Comparisons of the sex effects (with female as the reference) on individual protein abundance for (g) CC and DO, (h) CC and founder strains, and (i) DO and founder strains. N.S. indicates proteins that did not have significant sex effects. Sex effects are even more strongly correlated than heritability (*r* > 0.81, *p* < 2.2e-16). Red diagonal lines included for reference. A breakdown of the number of proteins with significant sex effects and their direction is shown for each comparison of populations.

## RESULTS

### Genetic effects on proteins are shared across the CC, DO, and founder strains

We quantified the abundance of 6,779 proteins in liver tissue from 58 inbred CC strains, one female and male per strain. We previously reported quantification of proteins from liver tissue of 192 outbred DO mice and 32 mice representing the eight founder strains (two per sex per strain) (Chick et al., 2016). The data for DO and founder strains were re-analyzed for this study to ensure that all data were processed consistently (**Methods**), resulting in the quantification of 6,588 and 6,950 proteins, respectively. From the 9,226 total proteins detected, 4,434 were observed in all three populations (**Figure 1b**; **Table S1**).

We estimated protein abundance heritability (*h^2^*), which reflects the combined effects of genetic variants, their allele frequencies, and the large-scale genetic background (*e.g.*, inbred or outbred) of each population (**Figure 1c**). Heritability estimates were highest in the CC and founder strains, due to capturing variation from non-additive genetic effects (*i.e.*, recessive and epistatic) in inbred strains. Heritability estimates in the DO are limited to additive genetic factors (narrow sense heritability; Lynch and Walsh 1998) that are captured in the genetic relatedness of our sample. Heritability estimates were significantly correlated across populations (*r* > 0.4, *p* <2.2e-16), suggesting that much of the controlling genetic variation is conserved across populations.

In order to identify the genetic loci that drive variation in protein abundance, we carried out protein quantitative trait locus (pQTL) mapping in the CC and DO (**Table S2**). To determine significant pQTL, we first applied a permutation analysis (Doerge and Churchill, 1996) to control genome-wide error rate for each protein and then applied a false discovery rate adjustment (FDR < 0.1) across proteins (Chesler et al., 2005) to establish a stringent detection threshold for pQTL. Using this criterion, we identified 1,199 local and 289 distal pQTL in the CC and 1,652 local and 382 distal pQTL in the DO (**Figure 2a & b**), where local is defined as when the pQTL is located within 10 Mbp of the midpoint of the protein-coding gene. We also identified a local pQTL on the mitochondrial genome in the CC for *mt-Nd1* (**Figure S1d**). In order to prevent the exclusion of true pQTL due to overly stringent control of false positive and accurately compare pQTL discovery across populations, we carried out a parallel analysis with more lenient FDR control (FDR < 0.5; **Figure S1a & b**).

**Figure 2.**
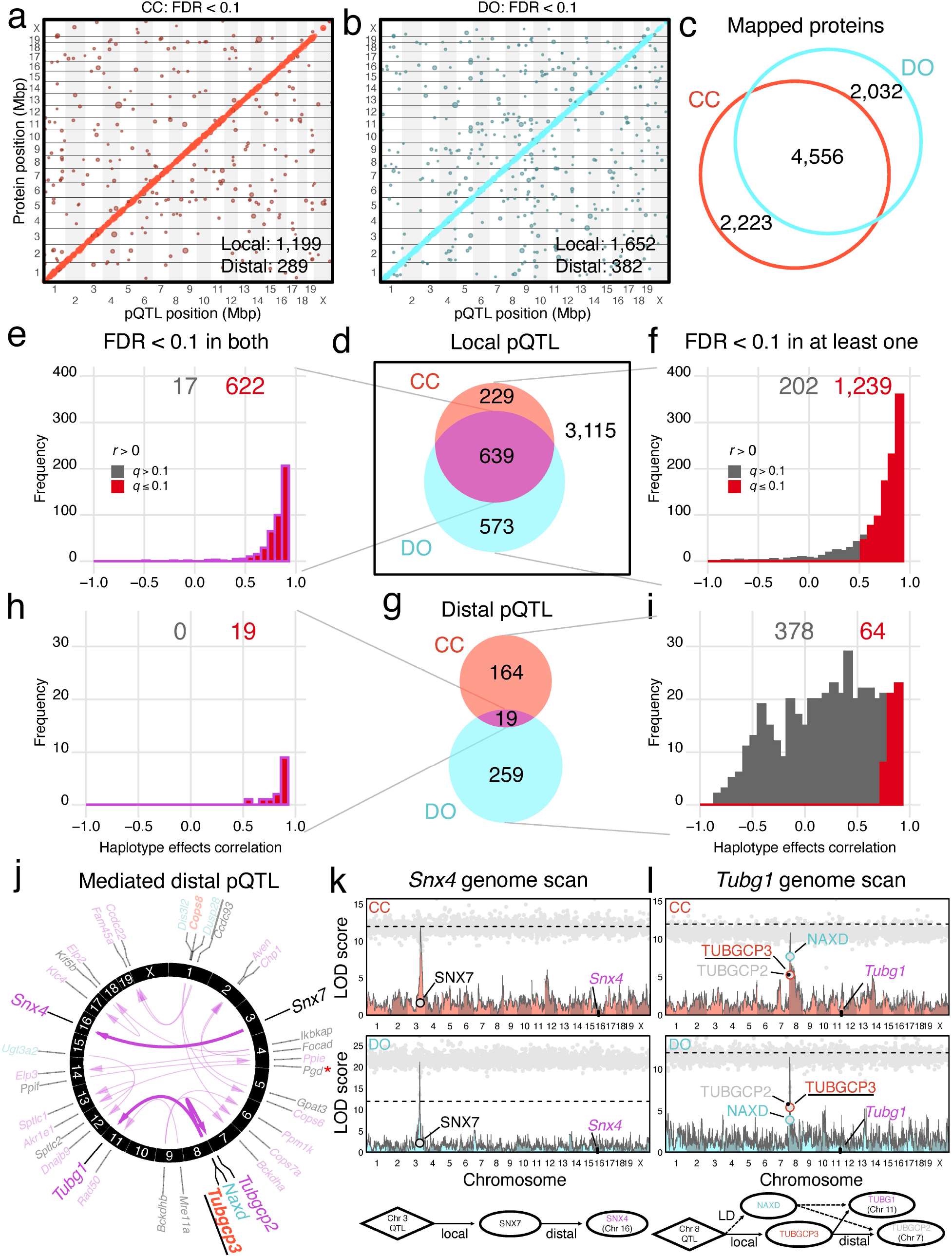
Genetic effects of loci are highly consistent between the CC and DO. Stringently detected pQTL (FDR < 0.1) in the (a) CC and (b) DO. The pQTL are plotted by the genomic positions of proteins against pQTL coordinates. Dot size is proportional to strength of association (LOD score). (c) Venn diagram of overlap of analyzed proteins for the CC and DO. (d) Venn diagram of overlap of local pQTL detected in the CC and DO. (e) The haplotype effects of local pQTL detected in both populations are highly consistent, as measured by the correlation coefficient comparing the effects in the CC and DO. (f) More local pQTL have consistent effects between the populations when also considering pQTL detected in only one of them. Red bars represent the number of pQTL that had significantly correlated effects (FDR < 0.1). (g) Venn diagram of overlap of distal pQTL detected in the CC and DO. (h) All 19 distal pQTL detected in both populations have consistent haplotype effects. (i) Considering distal pQTL detected in either of the populations resulted in 64 with correlated effects in the other population. (j) Circos plot of the 19 distal pQTL detected in both CC and DO. Arrows connect candidate drivers identified through mediation analysis to their targets (proteins with distal pQTL). Gene names in black represent the top candidates identified in both the CC and DO, whereas red and blue gene names were specific to the CC or DO, respectively. For candidate mediators that do not match between the CC and DO, the stronger candidate based on shared membership in protein families is underlined. The red asterisk denotes *Pgd* as a likely false positive protein mediator, observed in both the CC and DO. Examples of mediation agreement (*Snx4*) and disagreement (*Tubg1*) between the CC and DO are highlighted, with overlayed pQTL and mediation scans. Gray dots represent mediation scores for individual proteins. A strong mediator is local to the pQTL and has a steep drop from the peak LOD score. (k) SNX7 was detected as the driver of the *Snx4* distal pQTL in both the CC and DO. (l) TUBGCP3 was identified as the best candidate driver for the *Tubg1* distal pQTL in the CC, and NAXD as the third-best mediator; however, in the DO, NAXD was the stronger mediator. *Tubgcp3*, as well as *Tubg1* and *Tubgcp2*, are members of the tubulin superfamily, providing further support for TUBGCP3 as the stronger candidate driver of the distal pQTL of *Tubg1* and *Tubgcp2*. Horizontal dashed lines at LOD of 12 included for reference. Lenient mapping results (FDR < 0.5) further support the consistency of genetic regulation between the CC and DO (**Figure S1**).

We next restricted the set of proteins to the 4,556 that were detected and analyzed in both populations in order to compare genetic effects between the CC and DO. (**Figure 2c**). Among 1,441 local pQTL stringently detected for the shared proteins, 639 were detected in both populations (**Figure 2d**). To determine if these local pQTL were driven by the same genetic variants, we compared the estimated haplotype effects at each pQTL (**Methods**) and found that 622 (97.3%) were significantly positively correlated (**Figure 2e**; **Table S3**). To assess whether pQTL detected in only one population are population-specific or fell below the detection threshold in the other population but are likely driven by the same genetic variation, we compared the haplotype effects of detected pQTL to effects estimated at the corresponding locus in the other population, finding that 1,239 (86.0%) were significantly positively correlated. This trend for local pQTL holds true for lenient detection (**Figure S1f & g**; **Table S3**), indicating that local genetic effects on proteins are highly conserved between the CC and DO populations, even when failing to pass the threshold of detection in one of the populations.

The founder strains provide additional support for local pQTL in the CC and DO (**Figure S1k-n**), particularly for those that are challenging to detect due to rare alleles in the CC or DO (*e.g.*, a founder allele observed in three or fewer CC strains). We selected all genes with rare local founder alleles that did not have a leniently detected pQTL in the CC, representing 2,410 genes. We correlated the haplotype effects estimated at the locus closest to the gene transcription start sites (TSS) with the protein abundance in the founders (**Methods**) and found significant positive correlation for 196 genes, such as *Cyp2j5*. The conservation of local genetic effects among the founder strains, CC, and DO can be striking (such as in *Gosr2* and *Cyp2d22*; **Figure S2**). The three populations together identify a more complete catalog of local pQTL than any one alone. In total we find evidence to support local genetic effects on abundance for 3,004 proteins observed across the CC and DO.

Of the 183 distal pQTL stringently detected in the CC and 278 in the DO for the shared set of proteins, we found 19 that were stringently detected in both populations. All 19 also had significantly correlated haplotype effects (**Figure 2h**; **Table S3**). Overall, the distal pQTL are weaker than the local pQTL (*e.g.*, Chick *et al.* 2016; Albert *et al.* 2018) which may contribute to an increased rate of false negative findings. By comparing haplotype effects for pQTL that were only stringently detected in only one population, we identified an additional 45 shared distal pQTL between the CC and DO populations (**Figure 2i**; **Table S3**).

The genetic variants that drive distal pQTL are generally thought to act through diffusible intermediates, one or more of which are under local genetic control by the same variants. We used mediation analysis (Baron and Kenny 1986; MacKinnon *et al.* 2007), adapted for the DO (Chick *et al.* 2016; Keller *et al.* 2018) and CC (Keele et al., 2020), to identify candidate drivers of distal pQTL (**Methods**; **Table S4**). Mediation analysis can provide support for one or more candidate mediators for a pQTL. Furthermore, alternative candidates cannot be ruled out if they unobserved, such as proteins that failed to be detected by mass-spec or functional non-coding RNA. We identified candidate mediators for each of the 19 shared distal pQTL (**Figure 2j**). The same mediator was identified in the CC and DO for 14 of these – we note that BCKDHB, CCDC93, and IKBKAP are each candidate mediators of the distal pQTL for two proteins (CCDC93 for *Ccdc22* and *Fam45a*, BCKDHB for *Bckdha* and *Ppm1k* for, and IKBKAP for *Elp2* and *Elp3*). For four of the 19 distal pQTL, the best candidate mediator was different for the CC and DO. For example, TUBGCP3 is the strongest mediator of distal pQTL of *Tubg1* and *Tubgcp2* in the CC but NAXD is the strongest mediator in the DO (**Figure 2l**). Given that *Tubg1*, *Tubgcp2*, and *Tubgcp3* are all members of the tubulin superfamily, TUBGCP3 is a stronger candidate based on functional overlap. A true mediator should possess a local pQTL that co-localizes with the distal pQTL. Notably, the local pQTL for *Naxd* is much stronger in comparison to the local pQTL of *Tubgcp3* in the DO (LOD_*Naxd*_ = 40.2 compared to LOD_*Tubgcp3*_ = 11.0) whereas in the CC, they are more comparable (LOD_*Naxd*_ = 9.7 compared to LOD_*Tubgcp3*_ = 7.9). The local pQTL of *Naxd* in the DO may be acting as a surrogate for the local genotype, and thus outperforming TUBGCP3.

Similarly, COPS8 is the strongest mediator for distal pQTL of *Cops6* and *Cops7a* in the CC, whereas in the DO, COPS8 was not detected as a mediator and notably did not have a local pQTL (even leniently detected) in the DO, suggesting that COPS8 may be less accurately measured in our DO sample population. The final case of discordance was for the distal pQTL of *Dnajb9*, for which UGT3A2 was identified as a mediator in the DO whereas no candidate was found in the CC. *Ugt3a2* possesses local pQTL in both the CC and DO, which is notably strong in the DO (LOD_*Ugt3a2*_ = 36.9), suggesting it may be a false positive mediator in the DO and the true mediator was unobserved for both populations. We considered all distal pQTL that were stringently detected (FDR < 0.1) in one of the populations, and evaluated the corresponding pQTL status (stringent, lenient, or not detected) and mediation status (*e.g.*, same or different mediator) in the other population (**Figure S1o**), which revealed similar levels of concordance in mediation for pQTL detected in only one of the populations.

When we expand our pQTL comparisons to include all leniently detected pQTL, 1,462 proteins have significantly correlated local pQTL (223 more than with stringent detection in one population; **Figure S1g**). In contrast, fewer distal pQTL with significantly correlated effects are detected (22 compared to 64) (**Figure S1j**). Some pQTL that fail to meet the stringent threshold in either population still replicate across the CC and DO, as is the case for 12 of the 41 proteins with distal pQTL and correlated haplotype effects (**Figure S1i**). One interesting example is *Ercc3* (**Figure S3**), which has distal pQTL near a region on chromosome 7 that contains *Gtf2h1* and *Ercc2*, which all strongly associate with each other based on protein-protein interactions (Bioplex; Huttlin *et al.* 2020) and exhibit consistent but more extreme pQTL effects in the CC compared to the DO.

### Sex effects on protein abundance are extensively shared across populations

Protein abundances can differ between sexes (Chick et al., 2016), and this pattern can extend collectively to protein complexes (Romanov et al., 2019). We characterized sex effects in the CC, DO, and founder strains (**Methods**; **Table S1**). Significant sex effects (FDR < 0.1) were detected for 3,721 (54.9%) proteins in the CC, 4,376 (66.4%) proteins in the DO, and 1,549 (22.3%) proteins in the founder strains. The differences between male and female were overwhelmingly in the same direction for all populations (**Figure 1g-i**). Gene set enrichment analysis revealed that proteins related to ribosomes, translation, and protein transport gene ontology (GO) terms were more abundant in male livers whereas proteins related to catabolic and metabolic processes, including fatty acid metabolism, were more abundant in female livers in all populations.

### Drivers of variation in the abundance of protein-complexes

Members of protein-complexes exhibit varying degrees of co-abundance, with some groups of proteins being tightly co-abundant, *i.e.*, correlated, and others less so (Romanov et al., 2019). Correlation between members of a complex suggests some degree of co-regulation. We found that individual protein members of annotated protein-complexes (Giurgiu et al., 2019; Ori et al., 2016; Vinayagam et al., 2013; **Table S5**) are less heritable and fewer of them possess pQTL than proteins that are not members of complexes (**Figure S4a-d**; **Table S6**). However, protein-complexes can be influenced by genetic variation, as we previously reported for the Chaperonin containing T (CCT)-complex that is stoichiometrically regulated in response to the low abundance of a member protein, CCT6A (Chick et al., 2016). We evaluated the extent to which members of annotated protein-complexes were internally correlated, as well as how genetic factors and sex contribute to variation in their joint abundance. We summarized co-abundance, which we refer to as complex cohesiveness, as the median pairwise correlation between complex members (Romanov et al., 2019). To assess the contributions from genetic factors and sex, we summarized each protein-complex using the PC1 from member proteins. We first filtered proteins with local pQTL (FDR < 0.5) or strong distal pQTL (FDR < 0.1) to minimize the influence of individual proteins with independent genetic effects in order to focus on the shared genetic effects on a protein-complex. We estimated heritability and the proportion of variation explained by sex for each complex-specific PC1 (**Table S7**).

Complex-cohesiveness was correlated between the CC and DO (*r* = 0.70, *p* < 2.2e-16) (**Figure 3a & b**), and within each population, correlated with complex-heritability: *r* = 0.33, *p* = 4.37e-5 in the CC and *r* = 0.17, *p* = 0.03 in the DO (**Figure S4e**), suggesting that cohesiveness does reflect genetic factors that control protein abundance at some level. Notably, complex-heritability is consistently higher in the CC than DO (119 out of 163 complexes; 73.0%) and uncorrelated with complex-heritability in the DO (*r* = 0.08, *p* = 0.33), in contrast to the heritability of individual proteins (*r* = 0.43, *p* < 2.2e-16) (**Figure 3c & d**). The greater consistency between cohesiveness and heritability of protein-complexes in the CC is consistent with variation in overall protein-complex levels being influenced by non-additive genetic effects. The complex-heritability in CC captures the greater similarity of strain replicates with each other compared to other CC strains. The proportion of variation explained by sex for protein-complex abundance was correlated between the CC and DO (*r* = 0.48, *p* < 1.25e-10; **Figure 3e-f**). Protein-complexes previously shown to be driven by sex, such as eIF2B (Romanov et al., 2019), were confirmed in both the CC and DO.

**Figure 3.**
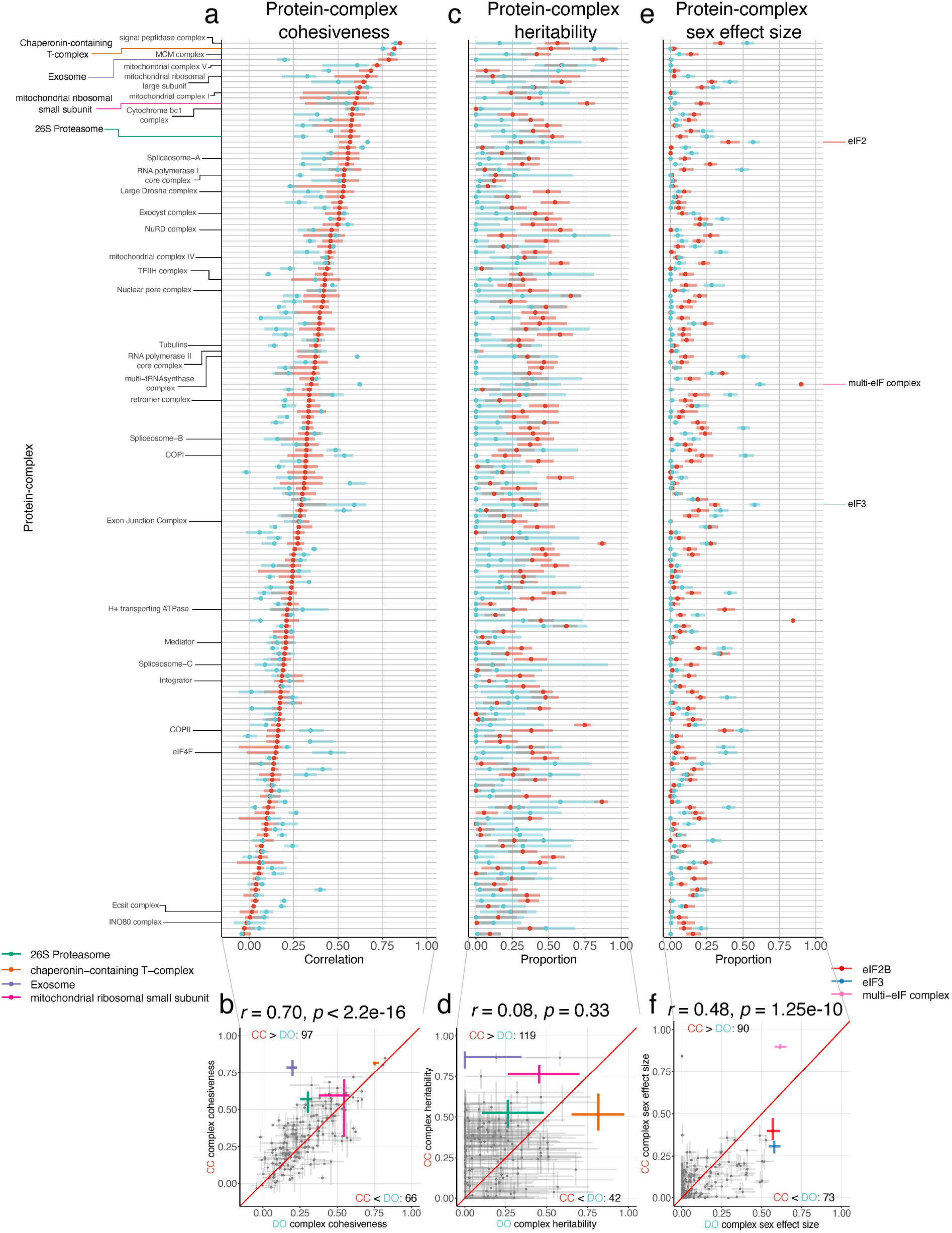
Genetic and sex effects on protein-complexes. (a-b) The co-abundance of complex members, *i.e.*, complex-cohesiveness, is consistent between the CC (red) and DO (blue) (*r* = 0.70, *p* < 2.2e-16). Cohesiveness is summarized (**Methods**) for each of 163 annotated protein complexes (Ori et al., 2016). (c-d) Complex-heritability, based on the first principal component (PC1) from each complex (**Methods**), were inconsistent between the CC and DO (*r* = 0.08, *p* = 0.33), with heritability notably higher more often in the CC. (e-f) Complex-sex effect size, representing the proportion of variability in the complex PC1 explained by sex, was consistent between the CC and DO (*r* = 0.48, *p* = 1.25e-10). For complex-cohesiveness, the intervals represent the interquartile range, and points represent the median. For complex-heritability and complex-sex effect size, intervals represent 95% subsample intervals. Exosome, chaperonin-containing T-complex, 26S Proteasome, and the mitochondrial ribosomal small subunit are highlighted as examples of highly heritable protein-complexes that are examined in detail (**Figures 4**, **5**, **6**, **S5**, & **S6**). Multi-eIF complex, eIF2B, and eIF3 are highlighted as complexes with large sex effect sizes in both the CC and DO, consistent with a previous study (Romanov et al., 2019). Red diagonal line included for reference.

### Genetic and stoichiometric regulation of the exosome

We identified the exosome complex as a novel and striking example of a protein-complex with stochiometric genetic regulation in the CC (**Figure 4**). The exosome complex had the highest complex-heritability in the CC (87.0%, [80.0-90.5%]) while being essentially non-heritable in the DO (0.0%, [0.0-34.6%]). Low EXOSC7 abundance is driven by a local PWK allele in the CC, which is homozygous in seven CC strains, whereas in our DO cohort, there were no mice homozygous for the PWK allele (**Figure S5b**). Heterozygous carriers of the PWK allele in the DO mice exhibited no discernable effect, suggesting that the exosome complex is regulated by a recessive PWK allele at *Exosc7* (**Figure S5c**), which is consistent with founder PWK mice having low EXOSC7 abundance (**Figure S5d**). Mediation analysis indicates that genetic variation at the *Exosc7* locus distally regulates the complex as well as the functionally related genes, *Dis3L* and *Etf1*. These two genes were not included in the original complex annotations, and their co-regulation suggests new biological interactions discovered via our approach. The complex-heritability (with *Dis3l* and *Etf1* included as well as complex members previously filtered out due to possessing pQTL) was 91.3%, and after removing the seven strains with the PWK allele at *Exosc7*, was reduced but still high at 61.4%, indicating the presence of additional genetic factors that affect the abundance of the exosome.

**Figure 4.**
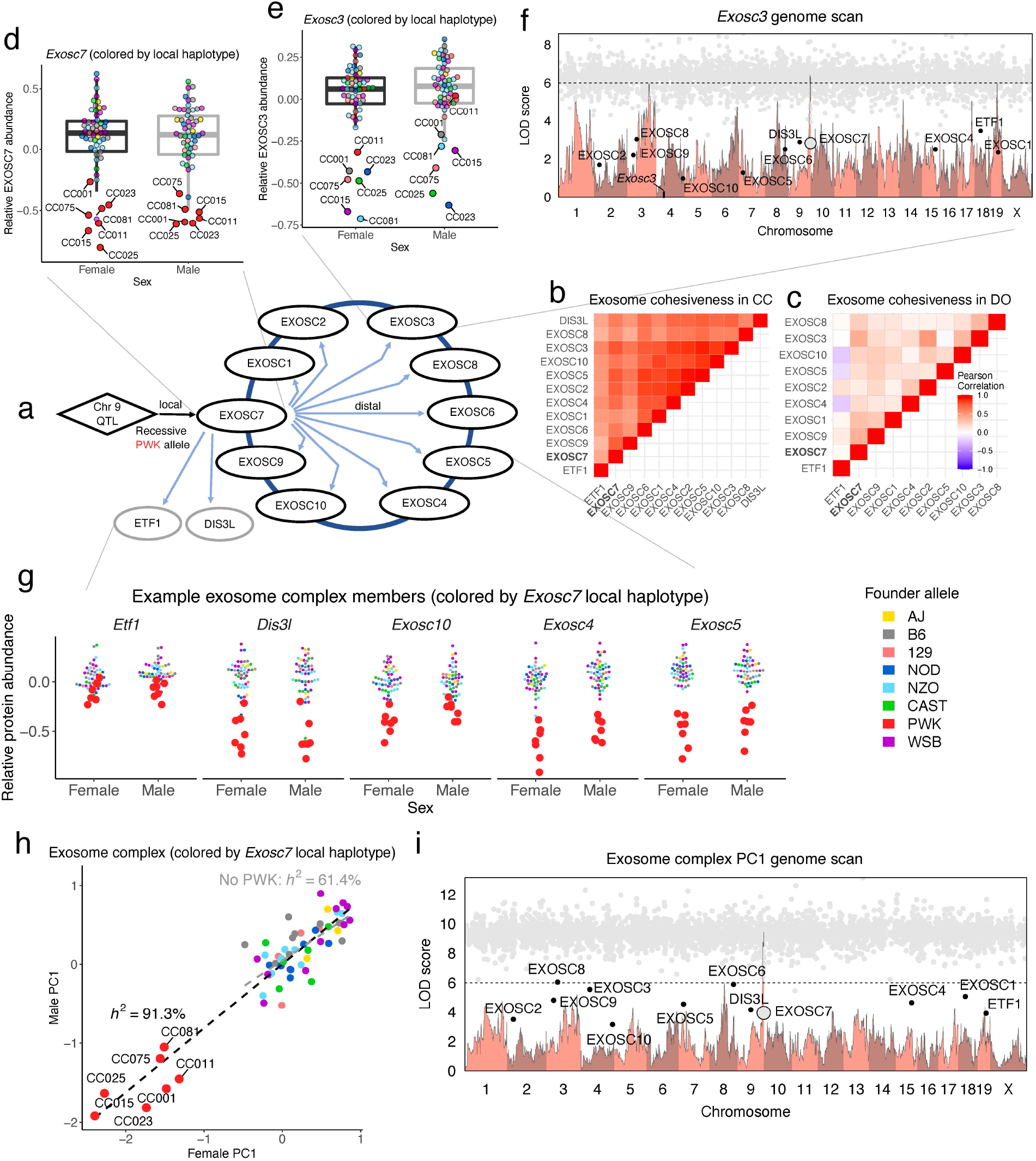
PWK allele at *Exosc7* drives low abundance of the exosome complex and related proteins. (a) Genetic variation at *Exosc7* has strong effects on the other members of the exosome complex and *Dis3l* and *Etf1*, which were not annotated members (Ori et al. 2016) though functionally related. The complex proteins were more tightly correlated in the (b) CC than the (c) DO, potentially due to the observance of the homozygous PWK genotype and strain replicates in the CC (**Figure S5**). (d) A local PWK allele resulted in lower EXOSC7 abundance, as observed in seven CC strains (CC001, CC011, CC015, CC023, CC25, CC075, and CC081). Points are colored by the haplotype allele at *Exosc7*. (e) Local genetic variation at *Exosc3* does not explain protein abundance patterns, which instead match the genetic variation at *Exosc7*, representing a distal pQTL. (f) The genome scan for *Exosc3* reveals the distal pQTL near *Exosc7*, and mediation analysis identifies EXOSC7 as a strong mediator of the distal pQTL (large gray dot) as well as the other exosome-related proteins as mediators (black dots), reflecting shared genetic regulation. The remaining smaller gray dots represent mediation scores for all other quantified proteins. Horizontal dashed line at LOD of 6 included for reference. (g) The PWK allele at *Exosc7* distally control other members of the exosome complex, with *Etf1*, *Dis3l*, *Exosc10*, *Exosc4*, and *Exosc5* shown here. Points are colored by the founder haplotype at *Exosc7*. (h) The first principal component (PC1) from the exosome complex, plotted males against females for the CC strains. Points are colored by the founder haplotype at *Exosc7*. CC strains separate based on whether they possess the PWK allele of *Exosc7*, reflected in a complex-heritability of 91.3%. After removal of the seven strains that possess the PWK allele, the complex-heritability is 61.4%, which suggest there are remaining loci that affect the abundance of the exosome. The black dashed line is the best fit line between males and females for the complex PC1, based on all 58 CC strains. The gray dashed line excludes the seven CC strains with the PWK allele at *Exosc7*. (i) The genome scan of the complex PC1 reveals a similar scan to *Exosc3*, though the pQTL near *Exosc7* and its mediation are stronger, due to the consistent effects across complex members.

### Secondary genetic effect on the chaperonin complex

Previously we found that the CCT complex was stoichiometrically regulated and driven by low abundance of CCT6A when the NOD haplotype is present (Chick et al., 2016). The CCT complex (**Figure 5**) has higher heritability in the DO (81.6%, [65.3-97.5%]) than in the CC (51.6%, [41.8-64.4%]) (**Figure 3c & d**). The DO sample is well-powered to detect the low NOD allele that drives the complex-wide regulation, due to 19 (9.9%) of the mice being homozygous NOD. The CC strains replicate the distal pQTL at the locus of *Cct6a* through a low NOD effect for complex members (*Cct4*, *Cct5*, *Cct8*, and *Tcp1*) – but CCT6a itself was not quantified in the CC samples.

**Figure 5.**
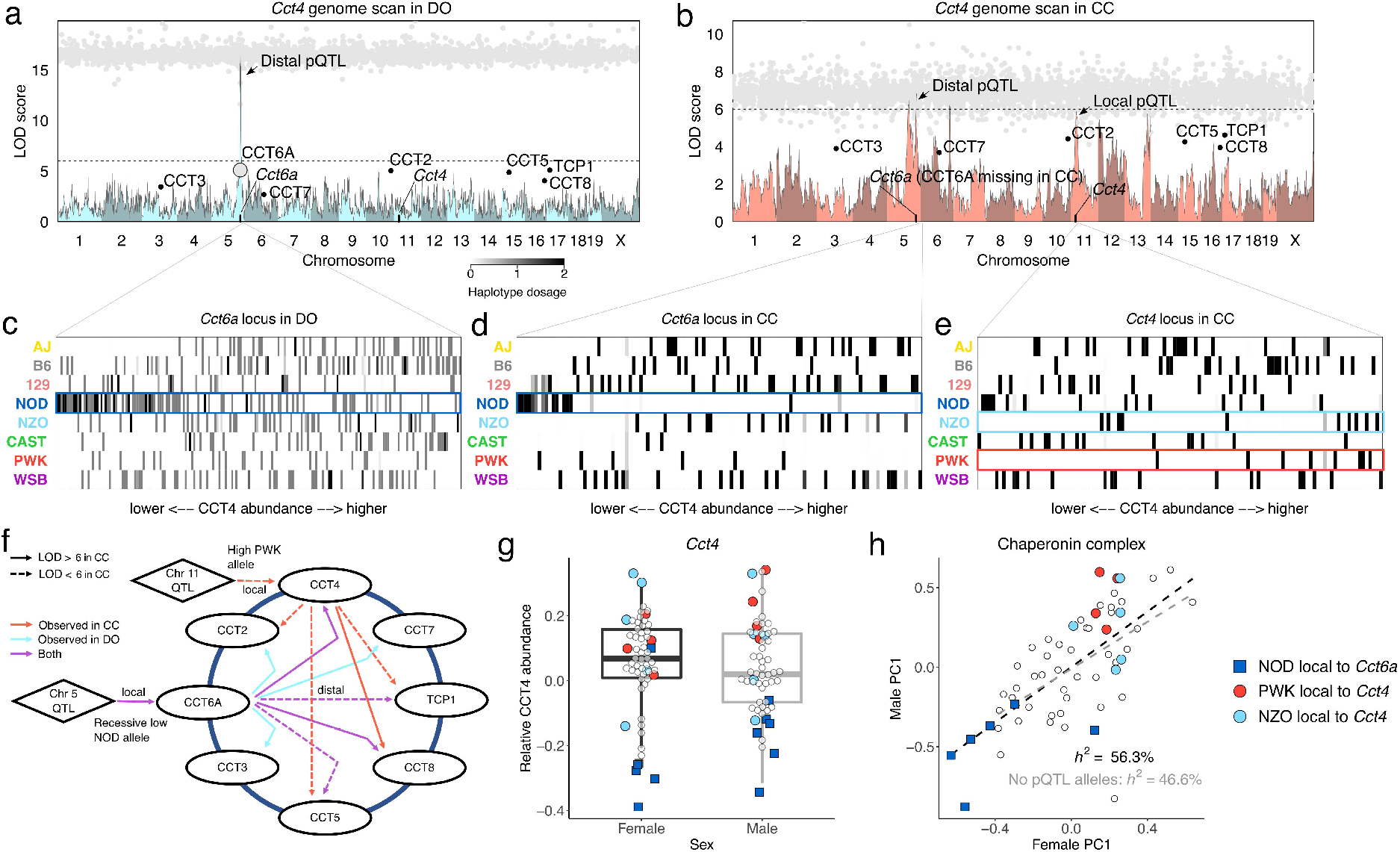
Genetic effects on the chaperonin-containing T-complex. Genome scans overlayed with mediation analysis for *Cct4*, a member of chaperonin-containing T (CCT)-complex, in the (a) DO and (b) CC, reveal a shared distal pQTL on chromosome 5. The CC also have a suggestive local pQTL for *Cct4* on chromosome 11. CCT4 was previously revealed through mediation analysis to be distally controlled through genetic variation at *Cct6a* (large gray dot) in the DO (Chick et al., 2016). The other CCT member proteins were detected as mediators (black dots) as well, stemming from shared genetic effects. The remaining smaller gray dots represent mediation scores of all other quantified proteins. The mediation scan of the distal pQTL on chromosome 5 in the CC similarly detects the other members of the CCT complex, though the likely causal intermediate CCT6A was not quantified in the CC. Horizontal dashed line at LOD of 6 included for reference. The founder haplotype inheritance, represented as heatmaps with founder allele dosage (expected allele counts) as rows and individual mice as columns, ordered by CCT4 abundance, at the chromosome 5 locus near Cct6a in the (c) DO and (d) CC and the (e) chromosome 11 locus near *Cct4* in the CC. A low NOD effect, potentially recessive and highlighted with dark blue boxes, drives the distal pQTL (likely through CCT6A) on CCT4 in both the DO and CC. The local effects on *Cct4* were not as strong, though NZO and PWK had subtle high effects, highlighted as light blue and red boxes. (f) A diagram of the population-specific and shared effects observed through pQTL and mediation analyses in the DO and CC. Many of the pQTL in the CC are weak, with all falling below the FDR < 0.1 threshold, and some with LOD < 6 (FDR > 0.5), but they are supported by overlap with the DO (chromosome 5) or being local to a complex member (chromosome 11). (g) In the CC, CCT4 abundance is affected by genetic variation at *Cct6a* as well as its local haplotype. The dark blue squares represent the six strains that possess the NOD allele at *Cct6a*, which had low CCT4. The light blue and red circles are the strains with high CCT4 that possess NZO and PWK alleles at *Cct4*. (e) The first principal component (PC1) for the complex, plotted males against females, reveals the same pattern of low strains (NOD allele of *Cct6a*; dark blue squares) and the high strains (NZO and PWK alleles of *Cct4*; light blue and red circles, respectively). The complex-heritability was high at 56.3% and after removal of the strains possessing notable alleles from pQTL at *Cct6a* and *Cct4*, dropped to 46.6%. The black dashed line represents the best fit line between males and females for complex PC1 based on all strains, and the gray dashed line is the best fit line with strains with the NOD allele of *Cct6a* and NZO and PWK alleles of *Cct4* excluded.

The effect of the pQTL at *Cct6a* drives less of the overall variation in the CC sample due to fewer occurrences of NOD homozygotes (12 CC mice in comparison to 19 DO mice). The CC reveals a secondary genetic effect mediated through CCT4, with high NZO and PWK alleles. After including complex members that were previously filtered out for possessing pQTL, the complex-heritability was 56.3%, which, after excluding CC strains that are NOD at *Cct6a* and NZO or PWK at *Cct4*, was still significant at 46.6% (**Figure 5h**), indicating that, as with the exosome complex, additional genetic effects contribute to CCT complex abundance.

### Independent genetic effects on the components and subcomplexes of the 26S proteasome

The 26S proteasome is composed of a 20S proteasome catalytic core (PSMA and PSMB proteins), which in the constitutive form has the constitutive subunits, PSMB5, PSMB6, and PSMB7, and is capped by two 19S regulators (composed of the PSMC and PSMD proteins). The constitutive form can be modified into the immunoproteasome by replacing the respective constitutive subunits with the three immunoproteasome inducible subunits, PSMB8, PSMB9, and PSMB10, and the 19S regulators with the 11S regulators (made up of PSME proteins) (Marshall and Vierstra, 2019; **Figure 6a**). The immunoproteasome is a highly efficient form of the proteasome that is predominantly expressed in immune cells (Kimura et al., 2015).

**Figure 6.**
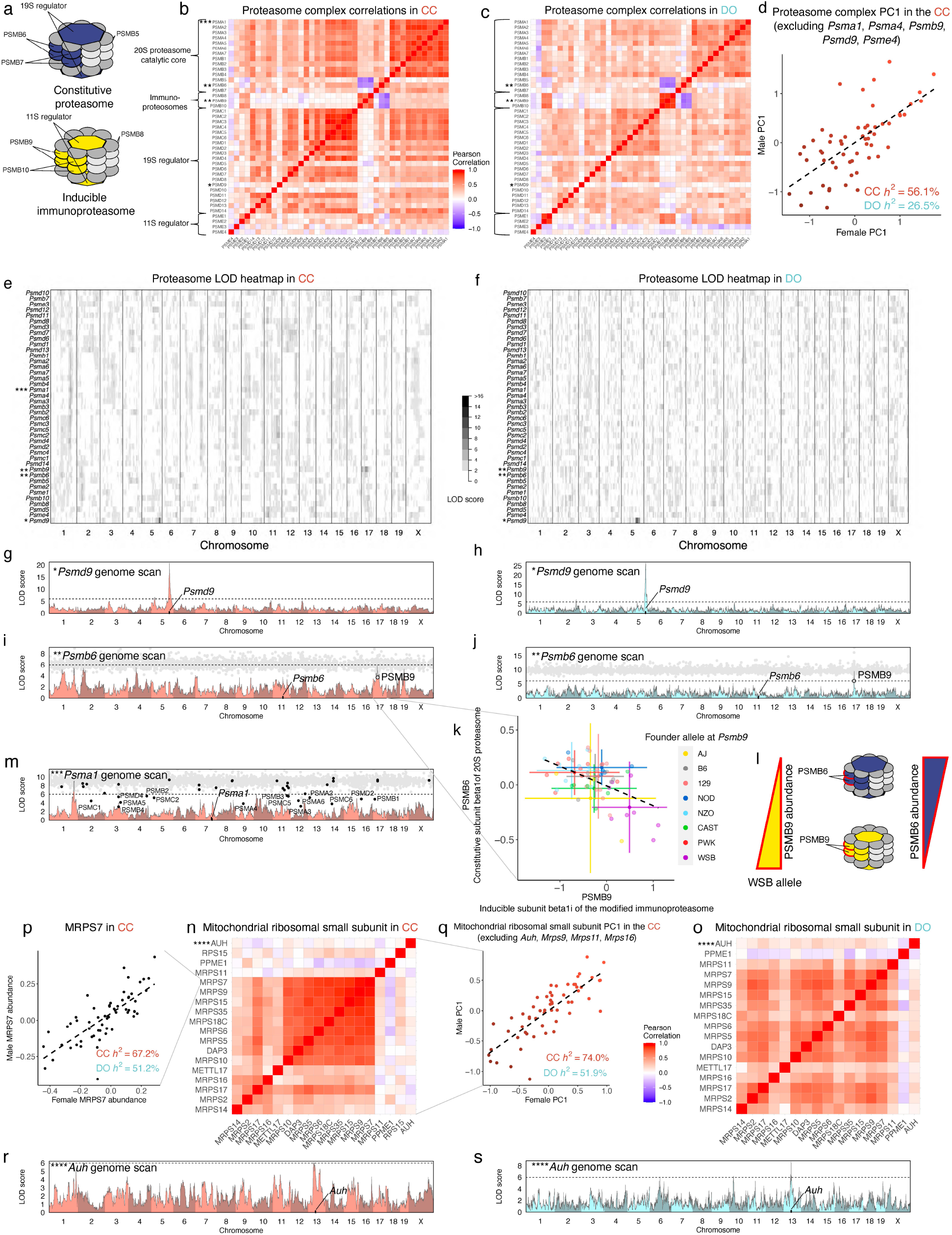
Polygenic regulation of the 26S proteasome and the mitochondrial ribosomal small subunit. (a) The 26S proteasome is composed of multiple subcomplexes: the 20S proteasome catalytic core (*Psma* and *Psmb* genes) and 19S regulator (*Psmc* and *Psmd* genes) for the constitutive form, and the inducible immunoproteasomes (*Psmb8*, *Psmb9*, and *Psmb10*) with their 11S regulator (*Psme* genes). The correlation patterns between member proteins reflect these subcomplexes, which are highly consistent between the (b) CC and (c) DO (and founder strains; **Figure S6a**). The overall correlation pattern reflects an inverse relationship between the constitutive proteasome and the inducible immunoproteasome. (d) CC strain identity explains a large portion of the variability in the first principal component (PC1) of the proteasome (after excluding genes with strong pQTL) and the complex-heritability is notably higher in the CC (56.1%) than DO (26.5%). The dashed line represents the best fit line between males and females in the complex PC1. A portion of the complex correlation structure can be explained by pQTL, shown here as a heatmap of genome scans (proteins as rows and genomic coordinate as columns), for the (e) CC and (f) DO. *Psmd9* (*) has a strong local pQTL in both the (g) CC and (h) DO, which is not shared with other proteasome proteins, explaining why *Psmd9* is poorly correlated with other members of the 19S proteasome. Horizontal dashed line at LOD of 6 included for reference. *Psmb6* (**) has a distal pQTL in both the (i) CC and (j) DO, which is mediated by PSMB9. Gray dots represent mediation scores for all quantified proteins, with PSMB9 highlighted as a large gray dot. (k) The relationship is negative – individuals with low abundance of PSMB9 (most notably CC strains with the WSB allele at *Psmb9*) have high abundance of PSMB6. Intervals represent mean ± 2 standard deviation bars. The dashed line represents the best fit line between PSMB6 and PSMB9 for the CC mice. (l) PSMB6 is a constitutive subunit of the 20S proteasome and PSMB9 is the corresponding inducible subunit in the modified immunoproteasome, suggesting that the inverse relationship between constitutive proteasome and immunoproteasome is genetically controlled for at least PSMB6 and PSMB9 in the CC and DO (though more broadly controlled in the founder strains; **Figure S6b-g**). A distal pQTL hotspot was observed in the CC on chromosome 1 (~170 Mbp) which was not present in the DO, for which (m) *Psma1* (***) has the strongest signal. Though a strong mediator was not detected from the proteins near the pQTL, many members of the 20S and 19S proteasomes present as strong mediators (black dots) due to the strong correlation among the proteins that map to the hotspot. The mitochondrial ribosomal small subunit was more heritable and cohesive than the 26S proteasome (**Figure 3**), which is evident in the correlation patterns for both the (n) CC and (o) DO. The individual proteins, such as (p) MRPS7, and the (q) complex PC1 (after excluding genes with strong pQTL) are highly consistent within CC strains despite few strong pQTL related to the complex. The dashed line represents the best fit line between males and females in MRPS7 and complex PC1. Similar to *Psmd9* within the 26S proteasome, a less cohesive member of the complex, *Auh* (****), can be explained by a unique local pQTL detected in both the (r) CC and (s) DO.

This alternation between two different forms of the proteasome is apparent in the anti-correlation between the inducible and immune components in both the CC and DO (**Figure 6b & c**) and founder strains (**Figure S6a**). The inverse relationship between the constitutive and inducible forms suggests that although individual samples vary in the relative abundance of the constitutive proteasome and immunoproteasome, they predominantly express one of the forms. Across the founder strains, this relationship appears to be genetically regulated, with the WSB, AJ, and NZO strains expressing more immunoproteasome, and the others expressing more of the constitutive form (**Figure S6b-g**). In the recombinant CC and DO, we identified a genetic variant that controls the balance between PSMB6 (constitutive) and PSMB9 (immunoproteasome). In both the CC and DO, genetic variation near *Psmb9* affected PSMB9 abundance, as well as PSMB6 abundance, confirmed through mediation analysis (**Figure 6i & j**). Consistent with the anti-correlation between inducible and constitutive subunits as well as the balance in the founder strains, the relationship between PSMB9 and PSMB6 is negative, *e.g.*, mice that inherited the WSB haplotype at the *Psmb9* locus have high PSMB9 abundance and low PSMB6 abundance (**Figure 6k & l**). The pQTL only explains the balance of PSMB6/PSMB9, and while consistent with the overall balance of the constitutive and inducible forms of the proteasome observed in the founder strains, it does not directly affect the other interchangeable members of the proteasome, which do not possess their own strong pQTL. The loss in the CC and DO of the co-regulation of all constitutive and inducible members suggests epistatic factors may interact with pQTL at *Psmb9* and those relationships are broken apart inthe recombinant populations, thus decoupling PSMB9 from the other immunoproteasome subunits.

Genetic factors also influence other components of the 26S proteasome. We identified a strong local pQTL that is consistent in both the CC and DO for *Psmd9* that does not affect other members of 19S regulator (**Figure 6g & h**), thus explaining the lack of cohesiveness of PSMD9 within the proteasome, which was previously observed in the DO (Romanov et al., 2019). A distal pQTL hotspot comprising members of the 20S catalytic core and 19S regulator mapped to a short interval on chromosome 1 in the CC, most strongly observed in *Psma1* (**Figure 6m**). No single, strong mediator was detected for the hotspot, though PEX19 was the best candidate for a number of the proteins, suggesting there may be multiple drivers of the hotspot or that the true driver was not observed at the protein-level.

### Polygenic regulation of the mitochondrial ribosomal small subunit

The mitochondrial ribosomal small subunit was highly cohesive in both populations (**Figures 3a, b**, **6n**, & **o**). The complex-heritability is also high, and after including all annotated members, was 74.0% in the CC and 51.9% in the DO. Despite high complex-heritability, we detected few pQTL for individual members of the complex. The one notable exception is *Auh*, for which we detected a strong local pQTL in the CC and DO (**Figure 6r & s**).

AUH was incohesive with the core of the complex, composed of mostly MRPS proteins. The complex PC1 and individual proteins are highly consistent within CC strain (*r* = 0.81 for the complex PC1; *r* = 0.76 for MRPS7) while displaying continuous variation across the CC population (**Figures 6p & q**). This distribution contrasts with the bimodal pattern of abundance for the exosome complex, which is driven by a single strong pQTL (**Figure 4h**) and suggests that many loci with small effects influence the overall abundance of the mitochondrial ribosomal small subunit.

### Strain-private variants affect protein abundance

Inbred mouse strains can accumulate mutations, and these variants can lead to phenotypic abnormalities across classical inbred strains, *e.g.*, new mutations that become fixed in sub-strains of B6 (Kumar et al., 2013) and in the parental B6 and DBA stocks (Anderson et al., 2002) used across different epochs of the BxD panel (Ashbrook et al., 2019; Mulligan et al., 2012).

New mutations originate and became fixed in the CC strains (Shorter et al., 2019; Srivastava et al., 2017) (**Figure 1a**). As a proof of concept, we first confirmed the loss of expression of proteins with known deletions, such as the 80 kbp deletion in CC026 that includes *C3* and a 15 bp deletion for CC042 in *Itgal* (**Figure 7a-b**) that increases susceptibility to tuberculosis (Smith et al., 2019) and salmonella (Zhang et al., 2019). Next, we estimated CC strain-specific effects for each protein in the CC and identified CC strains with extreme abundance of a given protein, referred to as strain-protein outliers (**Methods**). In total, we identified 6,046 strain-protein outliers, representing 4,323 proteins across all 58 CC strains. The strain-protein outliers coincided with 69 known strain-private genetic variants (Srivastava et al., 2017), meaning the strain-protein outlier could represent a local effect of the private variant associated with the protein’s coding gene, and this level of enrichment was significant per permutation (*p* = 3.7e-4). Interestingly, not all of the observed outliers associated with genetic variants were low extremes, as would be expected with a mutation that results in loss of the protein; we also observed increases in protein abundance in strains with private variants, such as *Sash1*, which harbors a novel SNP allele in CC058 (**Figure 7c**). CC004 has a unique SNP allele associated with low abundance of *Plek*, a gene which also has a weak local pQTL based on genetic variation from the founder strains (**Figure S7e & f**).

**Figure 7.**
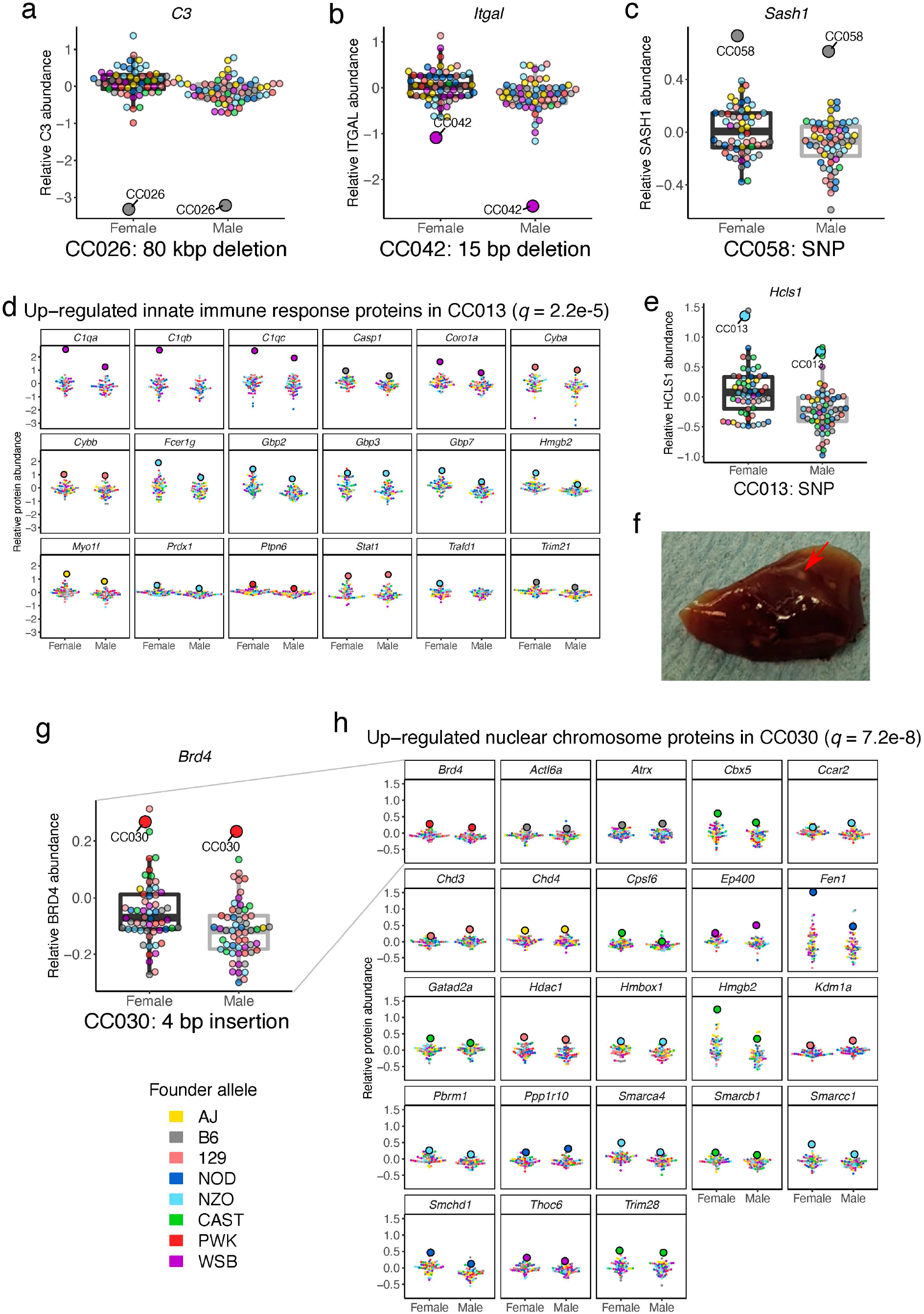
Strain-private genetic variants affect protein abundance and potentially influence larger protein networks. The effects of known strain-private deletions on protein abundance were confirmed for (a) *C3* in CC026 and (b) *Itgal* in CC042. The color or the dots indicates the founder allele at the gene. (c) A novel SNP allele in *Sash1* specific to CC058 is associated with an increase in SASH1 abundance. (d) Proteins related to innate immune response and other related pathways were high abundant in CC013. Large dots represent the specified CC strain with extreme protein abundance. (e) CC013 also possesses a strain-specific SNP allele in *Hcls1*, a gene involved in leukocyte differentiation, which may contribute to its unique protein abundance patterns in immune pathways. (f) CC013 has a unique liver phenotype, characterized by white granules, highlighted with red arrow, which may be related to the increased abundance of immune proteins. CC030 possesses a (g) private insertion in *Brd4*, which may contribute to (h) increased abundance in genes related to nuclear chromosome. Additional functionally related strain-specific protein dynamics are shown in **Figure S7**.

The presence of biologically related proteins with extreme abundances specific to CC strains further supports the impact of strain-specific genetic variants and regulatory patterns. We identified strain-specific dynamics by testing the set of outlying proteins specific to each CC strain for enrichment in GO and KEGG pathway terms (**Tables S8** and **S9**). In CC013, we observed increased abundance in proteins with GO annotations for the innate immune system (**Figure 7d**), leukocytes, and other immune system-related GO terms. CC013 possesses a unique SNP allele in *Hcls1* that was associated with increased HCLS1 abundance (**Figure 7e**), a gene involved in myeloid leukocyte differentiation that may contribute to the high abundance of immune-related proteins in CC013. CC013 also expressed a unique liver phenotype, characterized by white granules across the tissue (**Figure 7f**), which may relate to the excess of immune-related proteins. CC030 has a 4 bp insertion in *Brd4* that was associated with high BRD4 abundance and may contribute to increased abundance in proteins related to nuclear chromosome and chromatin (**Figure 7g-h**). The strain-specific protein outlier sets were enriched in a wide range of GO biological functions (**Figure S7g-j**), including low and high abundance of proteins from the mitochondrial respiratory complex I in CC007 and high abundance of cytosolic ribosome proteins in CC009.

## DISCUSSION

Multiparent populations are key resources for understanding genetic architecture; developing new phenotypic models of disease; and producing robust results that translate from model organisms to genetically diverse outbred populations such as humans. Here we show that the CC, DO, and their founder strains broadly share protein heritability, pQTL, their mediators, and sex effects on proteins. In comparison to individual proteins, protein-complexes were less consistent across these populations, highlighting the role of non-additive genetic effects in controlling protein-complex interactions. We observed a wide range of genetic effects on protein complexes, ranging from stoichiometric regulation in response to a large-effect locus (exosome and CCT complexes) to multi-locus and highly polygenic regulation (*e.g.*, the 26S proteasome and mitochondrial ribosomal small subunit, respectively). Within the CC, we observed strong effects on protein abundance from new mutations in specific CC strains. Lastly, we highlight individual CC strains with both aberrant protein regulation based on strain-specific mutations that broadly disrupt protein regulatory networks.

### Conservation of pQTL between CC and DO

Consistent with expectations, the effects of local genetic variation are highly conserved between the CC and DO (and often with the founder strains). In cases where discordance occurred between populations, we can often explain based on differing founder allele frequencies between the populations, such as for the local pQTL of *Ercc2* that is not detected in the DO because only one NOD homozygote was observed in the DO sample. Differences between populations stemming from different allele frequencies has been observed in human populations (Mogil et al., 2018). These discordant cases highlight how dominance and large-scale homozygosity contribute to the genetic effects on proteins, exposed by comparing the CC and DO. Distal (*i.e.*, *trans*) pQTL are often harder to detect due to weaker effects and are thus more likely to be discordant based on differences in allele frequencies and the presence of non-additive effects. Here we showed that the 19 distal pQTL that were detected in both populations at FDR < 0.1 are highly consistent, both in terms of haplotype effects and their mediation candidates. Discordance in effects between the CC and DO was much greater for distal pQTL detected in a single population, which is in striking contrast to the strong concordance of local genetic effects detected in only one population. The extent of discordance in distal pQTL effects compared to local effects suggests that additional factors are contributing beyond differing allele frequencies and may represent distal genetic effects that are not conserved across population.

Mediation analysis results mirror the concordance of distal pQTL effects, with largely the same mediators detected for distal pQTL detected with high confidence in both populations. There are some key caveats with mediation analysis, such as its accuracy being dependent on the true mediator being present in the data. If a candidate mediator is correlated with the true but unobserved mediator, possibly due to LD, it will likely be identified as a false positive mediator. This issue is problematic for proteomics studies if the mediator is not captured in the protein data, such as non-coding RNAs or lowly abundant proteins that go undetected in the mass-spec analysis. For example, PGD was detected as a candidate mediator of a distal pQTL of *Akr1e1* in both the CC and DO; however, previous studies identified zinc finger proteins (*Rex2* and *Zfp985*) as the most likely mediator of the pQTL (Hamilton-Williams et al., 2010; Keele et al., 2020). Zinc finger proteins are lowly expressed and not prevalent in proteomics data, and in their absence, the nearby protein PGD stands out as the best mediator in both populations. Variable measurement error across candidate mediators can also cause preference for a false mediator due to noise reducing the correlation between the distally controlled protein and its true mediator. This is likely occurring in the DO where NAXD is the stronger mediator for the *Tubg1* distal pQTL than TUBGCP3, the strong biological candidate. Furthermore, the presence of variable measurement error in proteomics is likely, stemming from the overall magnitude of each protein’s expression, the size of the protein and the number of peptides used to summarize it, and how specifically those peptides map to the protein. Comparisons of mediation analysis of two independent genetic experiments can provide replication of findings, as well as assess a more complete set of candidate mediators and correct misidentifications.

### Non-additive genetic effects on proteins and protein-complexes

Protein-complexes can be viewed as emergent phenotypes that can be driven by independently regulated members or sub-complexes, as well as higher order dynamics like protein-protein stoichiometry that control complex assembly. We observed a spectrum of cohesiveness (Romanov et al., 2019) across the protein-complexes and our three populations. These differences reflected components and sub-complexes that are semi-independent of the greater complex, as well as the underlying genetic architectures of the populations. Because high cohesiveness does not necessarily imply that a protein-complex is genetically regulated (*e.g.*, stoichiometry could produce tight correlation among complex members due to non-genetic factors), we employed genetic analyses (*e.g.*, QTL and heritability) to characterize genetic control of the protein-complexes. We found a number of strong and distinct examples of protein-complexes that are genetically regulated. Despite the challenges imposed by separate experiments and the relative nature of mass-spec proteomics (O’Brien et al., 2018), comparing the CC to the DO reveals the impact of an inbred genetic background on a number of protein-complexes, due to recessive effects, or conversely, the lack of dominance genome-wide.

Furthermore, CC strain replicates can capture multi-locus interactions, *i.e.*, epistasis, by fixing alleles at multiple loci within a strain. These results support the CC as a distinctly powerful tool for genetic analyses of emergent phenotypes like protein-complexes and as a companion resource to the DO for disentangling complicated genetic mechanisms.

The examples of genetically regulated protein-complexes that we identified represented a continuum of polygenicity underlying their genetic architecture. Examples ranged from the exosome with a single large effect pQTL local to a single member driving most variation, to the highly polygenic mitochondrial ribosomal small subunit, reflected in its continuous distribution across the CC strains and lack of pQTL. Intermediate to these extremes are examples like the CCT complex with its combination of a large effect pQTL with subtle secondary genetic effects revealed in the CC, and the 26S proteasome with its well-defined sub-complexes representing two distinct forms for altered biological function, the balance of which is genetically controlled to various degrees in these three populations. A common feature shared by these example protein-complexes examples is some degree of polygenicity. Even the fairly monogenic exosome and CCT complexes have some residual heritability after removing the CC strains that possess the contrasting alleles of the pQTL, suggesting other loci have secondary effects. The polygenic effects that drives the residual heritability may also be non-additive. In the DO, the exosome and CCT complex – after accounting for the strong pQTL of *Cct6a* – appear non-heritable, either due to large-scale dominance or the inability to capture epistatic effects in an outbred genetic background. These examples highlight the challenge of dissecting polygenic and non-additive effects down to individual loci and their specific mechanisms, emphasizing the value of related genetic resource populations with differing genetic architecture, like the CC and DO.

### New models of aberrant protein functional networks and disease

The unique biology of individual mouse strains has led to a variety of discoveries of genetically based disease phenotypes. We leveraged the CC population to identify strains exhibiting aberrant protein dynamics that could not be tied to the founder strains, such as being downstream of mutations unique to specific CC strains. They may also result from unique combinations of alleles of upstream drivers of a shared functional network. Examples include increased abundance of immune-related proteins and unique liver phenotype in CC013, which we are following up, and altered mitochondrial respiratory complex I function in CC007. Due to the replicability of the CC population, these strain-specific protein networks can be followed up, confirmed, and the underlying mechanisms dissected to reveal new biology and develop new models of disease.

### Integrative genetic resource populations

Proteins are a more functionally relevant measure of physiology and disease than the more commonly measured gene transcripts. In our examination of protein regulation across three related populations, we found that local genetic and sex effects on protein abundance were consistent, suggesting that molecular phenotype data and their findings can largely be integrated across these populations. It is easy to conceive of investigators querying specific genes of interest to assess whether their proteins possess sex effects, pQTL and their haplotype effects, or mediators of distal pQTL. To enable these queries, we provide interactive QTL analysis tools for both the CC (https://churchilllab.jax.org/qtlviewer/CC/Ferris) and DO (https://churchilllab.jax.org/qtlviewer/DO/Svenson). In contrast, higher order molecular phenotypes or characteristics, such as protein-complexes and regulatory networks, showed greater discordance across these populations. Non-additive genetic effects, either at a single locus or across loci, were apparent in the regulation of protein-complexes, driving the discordance between populations based on the genetic architecture of each. Lastly, we identified CC strain-specific aberrant protein abundances, their phenotypic or system relevance, and their putative consistency with previously described mutations present across these strains.

In this work, we used these diverse mouse populations to finely dissect genetic effects on protein-complexes in liver tissue, revealing the presence of non-additive effects and a polygenic spectrum of genetic regulatory patterns. In the future, we envision highly expandable joint population resources, covering a range of molecular phenotypes (*e.g.*, RNA-seq, ATAC-seq, mass-spec proteomics) and tissues relevant to human disease.

## METHODS

### Founder and CC strains

The CC and DO are descended from eight inbred founder strains: A/J (AJ), C57BL/6J (B6), 129S1/SvImJ (129), NOD/ShiLtJ (NOD), NZO/H1LtJ (NZO), CAST/EiJ (CAST), PWK/PhJ (PWK), and WSB/EiJ (WSB). We previously collected, processed, and quantified liver proteins from two females and two males from each founder strain as well as 192 DO mice (Chick et al., 2016).

We received pairs of young mice from 58 CC strains from the UNC Systems Genetics Core Facility between the summer of 2018 and early 2019. Mice were singly housed upon receipt until eight weeks of age. The 58 CC strains used in this study include: CC001/Unc (CC001), CC002/Unc (CC002), CC003/Unc (CC003), CC004/TauUnc (CC004), CC005/TauUnc (CC005), CC006/TauUnc (CC006), CC007/Unc (CC007), CC008/GeniUnc (CC008), CC009/UncJ (CC009), CC010/GeniUnc (CC010), CC011/Unc (CC011), CC012/GeniUnc (CC012), CC013/GeniUnc (CC013), CC015/Unc (CC015), CC016/GeniUnc (CC016), CC017/Unc (CC017), CC019/TauUnc (CC019), CC021/Unc (CC021), CC023/GeniUnc (CC023), CC024/GeniUnc (CC024), CC025/GeniUnc (CC025), CC026/GeniUnc (CC026), CC027/GeniUnc (CC027), CC029/Unc (CC029), CC030/GeniUnc (CC030), CC031/GeniUnc (CC031), CC032/GeniUnc (CC032), CC033/GeniUnc (CC033), CC035/Unc (CC035), CC036/Unc (CC036), CC037/TauUnc (CC037), CC038/GeniUnc (CC038), CC039/Unc (CC039), CC040/TauUnc (CC040), CC041/TauUnc (CC041), CC042/GeniUnc (CC042), CC043/GeniUnc (CC043), CC044/Unc (CC044), CC045/GeniUnc (CC045), CC046/Unc (CC046), CC049/TauUnc (CC049), CC051/TauUnc (CC051), CC053/Unc (CC053), CC055/TauUnc (CC055), CC057/Unc (CC057), CC058/Unc (CC058), CC059/TauUnc (CC059), CC060/Unc (CC060), CC061/GeniUnc (CC061), CC062/Unc (CC062), CC071/TauUnc (CC071), CC072/TauUnc (CC072), CC075/UncJ (CC075), CC078/TauUnc (CC078), CC079/TauUnc (CC079), CC080/TauUnc (CC080), CC081/Unc (CC081), and CC082/Unc (CC082). More information regarding the CC strains can be found at http://csbio.unc.edu/CCstatus/index.py?run=AvailableLines.information.

### Mouse genotyping, founder haplotype reconstruction, and gene annotations

The 116 CC mice were genotyped on the Mini Mouse Universal Genotyping Array (MiniMUGA), which includes 11,125 markers (Sigmon et al., 2020). Founder haplotypes were reconstructed using a Hidden Markov Model (HMM), implemented in the qtl2 R package (Broman et al., 2019), using the “risib8” option for an eight founder recombinant inbred panel. Notably, heterozygous markers are omitted, and haplotype reconstructions are limited to homozygous states, smoothing over potential residual heterozygous sites that remain in the CC mice. The genotyping and haplotype reconstruction for the DO mice were previously described (Chick et al., 2016); briefly, genotyping was performed on the larger MegaMUGA (57,973 markers) (Morgan and Welsh, 2015), from which founder haplotypes were reconstructed using the DOQTL R package (Gatti et al., 2014).

Ensembl version 91 gene and protein annotations were used in the CC, whereas version 75 was previously used in the older DO and founder strains data. If the gene symbol or gene ID differed for a protein ID between versions 75 and 91, we updated them to version 91 in the DO and founder strains. When comparing results (*e.g.*, pQTL, heritability, sex effects) between the CC and the DO or founder strains, we merged based on protein ID. For the more complicated results from mediation analysis, we allowed matches based on mediator gene symbol if the target protein IDs matched.

### Sample preparation for proteomics analysis

The sample preparation and mass-spec experimentation and analysis for the DO and founder strains were previously described (Chick et al., 2016). Singly housed CC mice had their food removed six hours prior to euthanasia and tissue harvest. Tissues were dissected out, weighed, and snap frozen in liquid nitrogen. Pulverized CC liver tissue were syringe-lysed in 8 M urea and 200 mM EPPS pH 8.5 with protease inhibitor and phosphatase inhibitor. BCA assay was performed to determine protein concentration of each sample. Samples were reduced in 5 mM TCEP, alkylated with 10 mM iodoacetamide, and quenched with 15 mM DTT. 200 μg protein was chloroform-methanol precipitated and re-suspended in 200 μL 200 mM EPPS pH 8.5. The proteins were digested by Lys-C at a 1:100 protease-to-peptide ratio overnight at room temperature with gentle shaking. Trypsin was used for further digestion for 6 hours at 37°C at the same ratio with Lys-C. After digestion, 50 μL of each sample were combined in a separate tube and used as the 11^th^ sample in all 12 tandem mass tag (TMT) 11plex. 100 μL of each sample were aliquoted, and 30 μL acetonitrile (ACN) was added into each sample to 30% final volume. 200 μg TMT reagent (126, 127N, 127C, 128N, 128C, 129N, 129C, 130N, 130C, 130N, and 131C) in 10 μL ACN was added to each sample. After 1 hour of labeling, 2 μL of each sample was combined, desalted, and analyzed using mass-spec. Total intensities were determined in each channel to calculate normalization factors. After quenching using 0.3% hydroxylamine, 11 samples were combined in 1:1 ratio of peptides based on normalization factors. The mixture was desalted by solid-phase extraction and fractionated with basic pH reversed phase (BPRP) high performance liquid chromatography (HPLC), collected onto a 96 well plate and combined for 24 fractions in total. Twelve fractions were desalted and analyzed by liquid chromatography-tandem mass spectrometry (LC-MS/MS).

### Off-line basic pH reversed-phase (BPRP) fractionation

We fractionated the pooled TMT-labeled peptide sample using BPRP HPLC (Wang et al., 2011). We used an Agilent 1200 pump equipped with a degasser and a photodiode array (PDA) detector. Peptides were subjected to a 50-min linear gradient from 5% to 35% acetonitrile in 10 mM ammonium bicarbonate pH 8 at a flow rate of 0.6 mL/min over an Agilent 300Extend C18 column (3.5 μm particles, 4.6 mm ID, and 220 mm in length). The peptide mixture was fractionated into a total of 96 fractions, which were consolidated into 24, from which 12 non-adjacent samples were analyzed (Paulo et al., 2016a). Samples were subsequently acidified with 1% formic acid and vacuum centrifuged to near dryness. Each consolidated fraction was desalted via StageTip, dried again via vacuum centrifugation, and reconstituted in 5% acetonitrile, 5% formic acid for LC-MS/MS processing.

### Liquid chromatography and tandem mass spectrometry

Mass spectrometric data were collected on an Orbitrap Fusion Lumos mass spectrometer coupled to a Proxeon NanoLC-1200 UHPLC. The 100 μm capillary column was packed with 35 cm of Accucore 50 resin (2.6 μm, 150Å; ThermoFisher Scientific). The scan sequence began with an MS1 spectrum (Orbitrap analysis, resolution 120,000, 350−1400 Th, automatic gain control (AGC) target 5E5, maximum injection time 50 ms). SPS-MS3 analysis was used to reduce ion interference (Gygi et al., 2019; Paulo et al., 2016b). The top 10 precursors were then selected for MS2/MS3 analysis. MS2 analysis consisted of collision-induced dissociation (CID), quadrupole ion trap analysis, automatic gain control (AGC) 1E4, NCE (normalized collision energy) 35, *q*-value < 0.25, maximum injection time 60 ms), and isolation window at 0.5.

Following acquisition of each MS2 spectrum, we collected an MS3 spectrum in which multiple MS2 fragment ions are captured in the MS3 precursor population using isolation waveforms with multiple frequency notches. MS3 precursors were fragmented by HCD and analyzed using the Orbitrap (NCE 65, AGC 3E5, maximum injection time 150 ms, resolution was 50,000 at 400 Th).

### Mass spectra data analysis

Mass spectra were processed using a Sequest-based pipeline (Huttlin et al., 2010). Spectra were converted to mzXML using a modified version of ReAdW.exe. Database search included all entries from an indexed Ensembl database version 90 (downloaded:10/09/2017). This database was concatenated with one composed of all protein sequences in the reversed order. Searches were performed using a 50 ppm precursor ion tolerance for total protein level analysis. The product ion tolerance was set to 0.9 Da. TMT tags on lysine residues, peptide N termini (+229.163 Da), and carbamidomethylation of cysteine residues (+57.021 Da) were set as static modifications, while oxidation of methionine residues (+15.995 Da) was set as a variable modification.

Peptide-spectrum matches (PSMs) were adjusted to FDR < 0.01 (Elias and Gygi, 2007, 2010). PSM filtering was performed using a linear discriminant analysis (LDA), as described previously (Huttlin et al., 2010), while considering the following parameters: XCorr, ∆Cn, missed cleavages, peptide length, charge state, and precursor mass accuracy. For TMT-based reporter ion quantitation, we extracted the summed signal-to-noise (S:N) ratio for each TMT channel and found the closest matching centroid to the expected mass of the TMT reporter ion. For protein-level comparisons, PSMs were identified, quantified, and collapsed to a peptide FDR < 0.01 and then collapsed further to a final protein-level FDR < 0.01, which resulted in a final peptide level FDR < 0.001. Moreover, protein assembly was guided by principles of parsimony to produce the smallest set of proteins necessary to account for all observed peptides. PSMs with poor quality, MS3 spectra with TMT reporter summed signal-to-noise of less than 100, or having no MS3 spectra were excluded from quantification (McAlister et al., 2012).

### Filtering out peptides that contain polymorphisms

Peptides that contain polymorphisms are problematic for protein quantification in genetically diverse samples because as the reference genome, only the B6 allele is quantified. Polymorphisms (with respect to the B6 genome) function as flags of the presence or absence of the B6 allele rather than reflecting the relative abundance of the peptide. During protein abundance estimation from peptides, a polymorphic peptide can either obscure the signal of a true abundance pQTL or induce a false local abundance pQTL. To avoid these biases, we filtered out peptides that contained polymorphisms based on the genome sequence of the founder strains and that were further confirmed in the data by having local founder haplotype effects that matched the expected distribution pattern of the B6 allele among the founder strains.

To determine whether peptides with polymorphisms matched their expected B6 allele distribution patterns, the peptide data was made more comparable by standardization within batches and removal of batch effects. Each peptide was scaled by a sample-specific within-batch scaling factor: 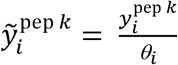, where 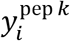 is the mass-spec intensity of peptide *k* for mouse *i*, 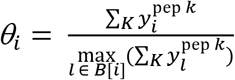, *K* is the set of all peptides measured for mouse *i*, and *B*[*i*] is the set of samples included in the mass-spec batch of mouse *i*. For the CC samples, the additional pooled sample (*i.e.*, bridge sample) was also included in each batch and provided an additional standardization across batches: 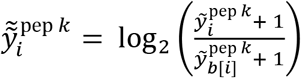, where *b*[*i*] represents the bridge sample from the batch of mouse *i*. For the DO and founder strain samples that did not include bridge samples, 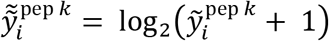. A log transformation is used because peptide intensities are commonly log-linear.

Batch effects were removed from the processed peptide data using a linear mixed effect model (LMM) fit with the lme4 R package (Bates et al., 2015). Peptides unobserved for all samples within a batch were recorded as missing (coded as NA). If greater than 80% of samples were missing for a polymorphic peptide, it was removed from the batch correction step and the subsequent evaluation of the B6 allele distribution pattern. The following model was fit for the CC data:

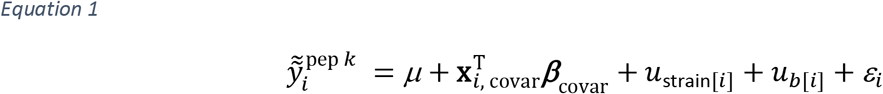

where *μ* is the intercept, ***β***_covar>_ is the effect vector of covariates estimated as fixed effects, 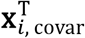 is the *i*^th^ row of the covariate design matrix, *u*_strain[*i*]_ is the effect of the strain of sample *i*, *u*_*b*[*i*]_ is the effect the batch of sample *i*, and *ε*_*i*_ is the error for sample *i* with *ε*_*i*_~N(0, σ^2^). The strain and batch effects were estimated as random effects: 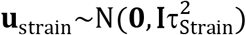 and 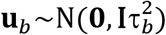. For the CC and founder strains, sex was included as a covariate. A similar model was fit for the DO but with no strain effect and diet also included as a covariate with sex. The batch effects, estimated with best linear unbiased predictors (BLUPs) using restricted maximum likelihood estimates (REML; Patterson and Thompson 1971), were subtracted from each peptide measurement: 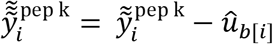.

For peptides expected to contain a polymorphism, we then fit local genetic effects: haplotype effects at the marker closest to the TSS of the gene to which the peptide maps in the CC and DO and strain effects in the founder strains.

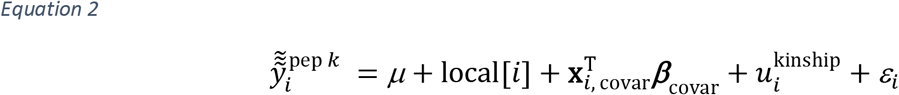

where local_*i*_ is the effect of the local haplotype pair to peptide *k* for sample *i*, 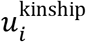 represents a random effect that accounts for the correlation structure between individuals that is consistent with overall genetic relatedness, often referred to as the kinship effect, and all other terms as previously defined. For the CC, 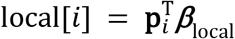 where 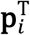 is the founder haplotype probability vector at the marker closest to the gene TSS (*e.g.*, ordering the founder strains as AJ, B6, 129, NOD, NZO, CAST, PWK, and WSB, 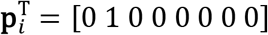 for a CC mouse *i* that is B6/B6 at the locus). For the DO, 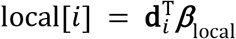 where 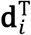 is the founder haplotype dosage vector, scaled to sum to zero, at the marker closest to the gene TSS (*e.g.*, 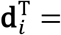 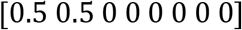 for a DO mouse *i* that is AJ/B6 at the locus). For the founder strains, 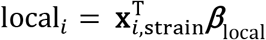 where 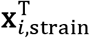 is the founder strain incidence vector for mouse *i* (*e.g.*, 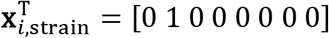 for a B6 mouse). ***β***_local_ is an eight-element vector of founder haplotype effects, fit as a random effect: 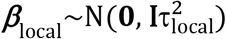 where **I** is an 8×8 identity matrix and 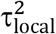 is the variance component underlying the local effects. The kinship effect is included for the CC and DO and modeled as 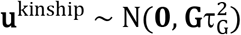 where **G** is a realized genomic relationship matrix and 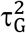 is the variance component underlying the kinship effect, accounting for population structure (Kang et al., 2008, 2010; Lippert et al., 2011; Zhou and Stephens, 2012). Here we used a leave-one-chromosome-out or “loco” **G**, in which markers from the chromosome the peptide is predicted to be located on are excluded from **G** estimation in order to avoid the kinship term absorbing some of local[*i*] (Wei and Xu, 2016). We then calculated 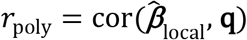, the Pearson correlation coefficient between 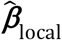, the BLUP of ***β***_local_ and **q**, the incidence vector of the B6 allele among the founder strains (*e.g.*, **q** = [0 1 0 0 0 0 0 0] for a peptide with a B6 allele that is missing in the other founder strains). Sets of peptides with polymorphisms were defined based on having *r*_poly_ > 0.5 for each of the CC, DO, and founder strains, to be excluded from further analysis because they would bias protein abundance estimation.

### Protein abundance estimation from peptides

Protein abundances were estimated from their component peptides after filtering out peptides that possessed polymorphisms based on founder strain sequences that were confirmed in the peptide data. The abundance for protein *j* is calculated as 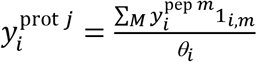 where *M* is the set of peptides that map to protein *j*, 1_*i,m*_ is the indicator function that peptide *m* was observed in mouse *i*, and *γ*_*i*_ is the scaling factor previously defined (Huttlin et al., 2010). Similar to the previously described peptide normalization in the CC, proteins were scaled relative to the bridge sample (pooled sample of all CC mice included in all batches) and log-transformed: 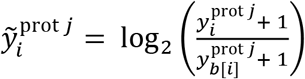. For the DO and founder strain samples, there was no bridge sample, and proteins were instead normalized as: 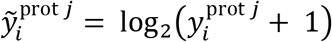.

Batch effects were removed from the protein data using the same LMM approach described for the peptide data (Equation 1). If more than 50% of samples were missing a protein, it was removed from further analysis. Batch effects, estimated as BLUPs, were then removed: 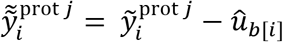.

### Heritability estimation

We estimated heritability for all proteins in the CC, DO, and founder strains. The heritability model is similar to Equation 2, but for proteins instead of peptides and without the local[*i*] term:

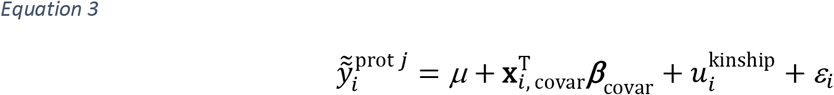

where terms are as previously defined. The genomic relationship matrix **G** – corresponding to the kinship term 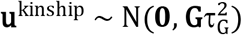 for the CC and DO – is estimated from all markers –“non-loco” **G** – because there are no other genetic terms in the model. In the founder strains, 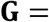 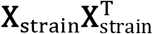 where **X**_strain_ is the founder strain incidence matrix. Sex was modeled as a covariate for all three populations, and diet as well in the DO. Heritability is then calculated as 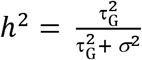

### QTL analysis

In the CC and DO, we performed a genome-wide pQTL scan for each protein, testing a QTL effect at positions across the genome, using a model similar to Equation 2:

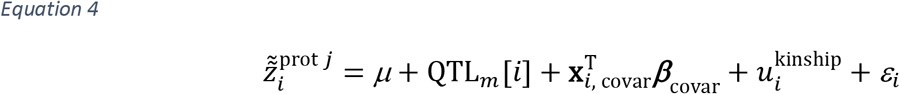

where 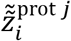 is the standard normal quantile returned by the inverse cumulative distribution function of the normal distribution on the uniform percentiles defined by the ranks of 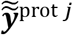, *i.e.*, the rank-based inverse normal transformation (RINT) (Beasley et al., 2009) of protein *j* for individual *i*, QTL_*m*_[*i*] is the effect of the putative QTL at marker *m* on protein *j* for individual *i*, equivalent to the local[*i*] term in Equation 2 for the CC and DO, and all other terms as previously defined. The kinship effect was fit based on the “loco” **G** specific to the chromosome of marker *m*. We used RINT for the QTL analysis to reduce the influence of extreme observations that can produce false positives, particularly when they coincide with a rare founder haplotype allele, which are of particular concern in the context of a CC sample population with 58 unique genomes. To test the QTL term, the model in Equation 4 is compared to a null model excluding QTL_*m*_, summarized as the log_10_ likelihood ratio (LOD) score.

The QTL model in Equation 4 was also used for variant association mapping at specific pQTL identified through the haplotype-based analysis by adjusting the QTL_*m*_[*i*] term: 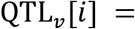 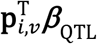, where 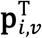 is the marginal variant allele probability vector for variant *v*, which is calculated by collapsing and simplifying the underlying founder haplotype probabilities based on the known variant genotype in the founder strains (SQLite variant database: https://doi.org/10.6084/m9.figshare.5280229.v3).

For the CC, we mapped pQTL based on strain averages where 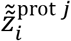 is the average of 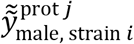 and 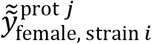 followed by RINT across the population of strains. Founder male, strain *i* female, strain *i* haplotype probabilities were reconstructed at the level of individual mice and were also averaged at all markers for strain-level mapping. A strain-level **G** was then estimated from the strain-level founder haplotype probabilities. No fixed effect covariates were included when mapping on strain averages. We tried mapping pQTL in the CC on individual-level data, which returned largely consistent results, but notably fewer and weaker pQTL. In the CC, we also mapped pQTL to the mitochondrial genome and Y chromosome by testing whether the founder origin of the mitochondria or Y chromosome was associated with protein abundance. We fit Equation 4, treating the mitochondrial genome or Y chromosome as a single locus QTL_Y_[*i*] and QTL_MT_[*i*], respectively, using the “non-loco” **G** for the kinship effect. The founder strain of origin for the Y chromosome was determined for all CC strains. For the mitochondrial genome, six strains (CC031, CC032, CC041, CC051, CC059, CC072) possessed ambiguity between AJ and NOD, which we encoded as equal probabilities 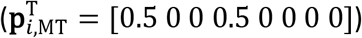.

### QTL significance thresholds

We estimated significance thresholds for pQTL using permutations (Doerge and Churchill, 1996). We accounted for varying levels of missing data by performing genome scans on 10,000 permutations of the normal quantiles for each level of observed missingness in the CC and DO (ranging from 0 to 50%). Genome scans of the permuted data used the model in Equation 4, while excluding covariates and the kinship term, allowing permutations to be more applicable across proteins with the same level of missingness. To control the genome-wide error rate per protein and the FDR across proteins (Chesler et al., 2005), the maximum LOD scores from the permutation scans were used to fit generalized extreme value distributions (GEV) (Dudbridge and Koeleman, 2004; Valdar et al., 2009) specific to the level of missingness, which were used to calculate genome-wide permutation *p*-values for the maximum LOD observed per protein:

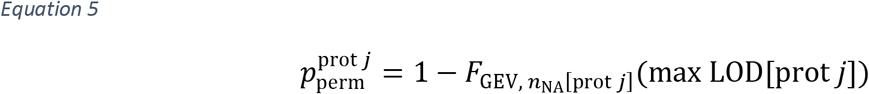

where 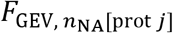 is the cumulative density function for the GEV fit from the permutations of quantiles with *n*_NA_ number missing values, corresponding to the number missing for protein *j*, and max LOD[prot *j*] is the maximum LOD score from the genome scan of protein *j*. We then used the Benjamini-Hochberg (BH) procedure (Benjamini and Hochberg, 1995) to calculate FDR <*q*-values from all observed permutation p-values, which were used to interpolate a permutation *p*-value that corresponds to 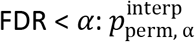 where α ∈ [0.1, 0.5]. Significance thresholds on the LOD scale, specific to FDR < α and *n*_NA_ missing data points, were calculated: 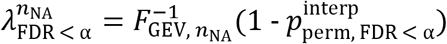 where 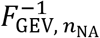 is the inverse cumulative density function for the GEV with *n*_NA_ missing data points. As a final step to reduce random variation between sets of permutations, we regressed the estimated thresholds for a population and FDR level on the number of missing data points *n*_NA_, and created a table of fitted thresholds: 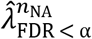 for α ∈ [0.1, 0.5] for both the CC and DO. Whether a pQTL met FDR < α significance, the threshold corresponding to α with the *n*_NA_ for protein *j* was used. For reference, 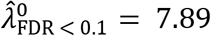 and 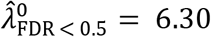 in the CC, and 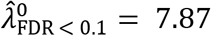 and 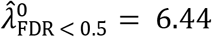 in the DO.

### Consistency of QTL between the CC and DO

We evaluated the consistency of pQTL between the CC and DO by comparing their haplotype effects. Haplotype effects were estimated at the pQTL marker using the model in Equation 4. To stabilize the effects, they were modeled as a random effect: 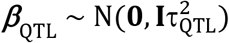, where 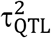 is a variance component underlying the haplotype effects of the pQTL. We then estimated the haplotype effects as BLUPs 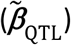. To declare pQTL consistent between the CC and DO, we evaluated whether their haplotype effects were significantly positively correlated: 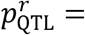 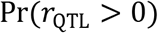 where 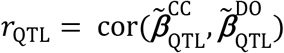 and 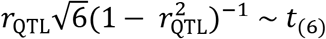. To account for multiple testing, we used the BH procedure on the *p*-values for correlated effects and declared pQTL with 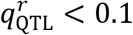 as consistent between the CC and DO.

Selecting which marker to fit the haplotype effects at is complicated by the fact that the CC and DO have different sets of markers and the genomic coordinate of the peak LOD score is also subject to variation. When comparing pQTL detected in both populations, we fit Equation 4 at the markers with the highest LOD score specific to each population, meaning they may not be the markers closest to each other. When comparing pQTL that were detected in only one population, we selected the marker in the population that failed to map the pQTL that was closest to the marker in the population that detected it.

### Consistency of local QTL in the CC with the founder strains

If the genetic effects on a protein are primarily local, the relative abundances for a protein in the founder strains should match the local pQTL effects observed in the CC and DO, which can be used to confirm findings and better support suggestive pQTL in the CC or DO. We evaluated the consistency of local pQTL in the CC with the founder strains, using an approach similar to how we compared pQTL effects between the CC and DO. For the founder strains, rather than fitting pQTL effects 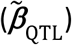, we fit the founder effects as random terms (as described for the local term in Equation 2 for the founder strains) summarized as BLUPs 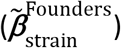. We then calculated the Pearson correlation between local pQTL effects in the CC and founder effects in the founder strains: 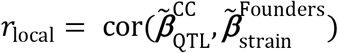. As when comparing QTL effects between the CC and DO, we then tested *r*_local_ > 0, and corrected for multiple testing through the BH procedure.

### Mediation analysis

We performed mediation analysis on each distal pQTL with LOD > 6 in the CC or DO, which involved a scan analogous to the QTL genome scans. The underlying model is

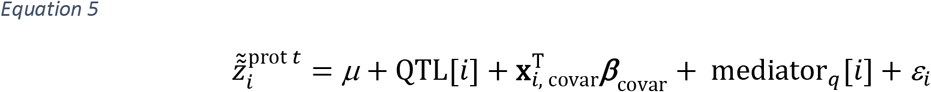

where QTL[*i*] is as defined for QTL_*m*_[*i*] in Equation 4, but fixed at the marker *m* of the detected distal pQTL for target protein *t*, mediator _*q*_[*i*], is the effect of candidate mediator protein *q* on protein *t* for individual *i*, and all other terms as previously defined. The effect of the mediator is modeled as: 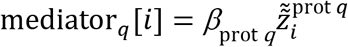, where *β*_prot *q*_ is the regression coefficient for the mediator protein *q* and 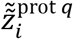 is the RINT quantity of protein *q* for individual *i*. The likelihood of Equation 5 is compared to a null QTL model that excludes the QTL_*i*_ term, producing a mediation LOD score. The mediation model is fit for all proteins as mediators, excluding protein *t*, resulting in a mediation scan.

We assume that the vast majority of candidates are not true mediators of a specific pQTL and thus the distribution of mediation LOD scores approximates a null distribution. Assuming that the null distribution is approximately normal, we calculate the z-scores of the mediation LOD scores, and define mediators of the QTL at marker *m* for protein *t* as proteins with 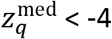, where 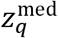 is the z-score of the mediation LOD score for candidate mediator protein *q*. The rationale being that when testing the QTL term in Equation 5, if the mediator contains some of the information of the QTL, its presence in both the alternative and null models will result in a large drop in the LOD score of the detected pQTL. In order for a mediator to be a clear candidate driver of the distal pQTL, we required that the mediator TSS be within 10 Mbp of the pQTL marker. Strong mediators that were not near the pQTL often represent proteins that are correlated with protein *t*, such as co-regulated members of a protein-complex.

### Sex effects on proteins analysis

Proteins that exhibited differential abundance between the sexes, *i.e.*, sex effects, were identified using an LMM similar to the heritability model (Equation 3) for the CC, DO, and founder strains, but instead testing the significance of the sex coefficient:

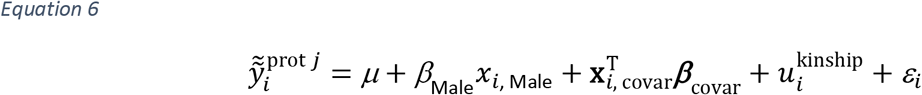

where *β*_Male_ is the effect on protein *j* of being male, *x*_*i*, Male_ is an indicator variable of being male, and all other terms as defined previously. Non-sex covariates and the specification of 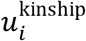 for the different populations are the same as described for heritability.

A *p*-value for the sex effect was calculated by comparing the model in Equation 6 to a null model without the sex effect through the likelihood ratio test 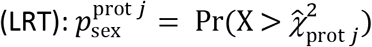 where Pr(.) denotes the 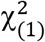 probability density function and 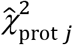 is the observed LRT statistic for protein *j*. The LMM was fit with the qtl2 R package (Broman et al., 2019), using maximum likelihood estimates (MLE) for parameters rather than REML, which are more appropriate for asymptotic-based significance testing of fixed effects. Proteins with significant sex effects were selected based on FDR < 0.1 using the BH procedure (Benjamini and Hochberg 1995).

We performed gene set enrichment analysis through the clusterProfiler R package (Yu et al., 2012). We defined gene sets based on having *q*_sex_ < 0.01 and split them further into subsets based on having higher abundance in males or higher abundance in females for both the CC and DO. We used the quantified proteins in each population as the background gene set to account for biases in the observed proteins. Hypergeometric set tests for enrichment of GO and KEGG terms were performed with FDR multiple testing control (Storey et al., 2019). Enriched GO and KEGG terms were selected based on having *q*_set_ < 0.1.

### Protein-complex analysis

We categorized individual proteins as members of protein-complexes as defined by previous annotations (Ori et al., 2016). For each protein-complex, we quantified how tightly co-abundant, *i.e.*, cohesive, the members are, by calculating the median pairwise Pearson correlation for each protein with the other members of the protein-complex. We summarized cohesiveness within a protein-complex by recording the median and interquartile range across the median correlations for the individual proteins.

To assess whether genetics or sex regulated protein-complexes as a whole, we estimated the complex-heritability and complex-sex effect size. We took the PC1 from the set of proteins from the complex after first removing the effects of key covariates from the individual proteins in order to keep the PC1 from reflecting other drivers of variation instead of genetic factors or sex. For complex-heritability, we removed the effect of sex in the CC, and both sex and diet in the DO. For complex-sex effect size, we only removed the effect of diet from the DO. Complex-heritability was estimated using the model in Equation 3, with no covariates and the complex PC1 as the outcome. To estimate the complex-sex effect size: 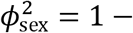 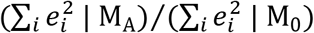 where 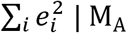 is the sum of squared residuals (SSR) under the alternative model (Equation 6) and 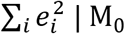 is the SSR under the null model (Equation 6 excluding sex effect). Interval estimates for complex-heritability and complex-sex effect size represent 95% subsample intervals. We randomly sampled without replacement 80 of the CC and DO data 1,000 times and estimated the complex-heritability and complex-sex effect size for each subsample as well as the 2.5^th^ and 97.5^th^ quantiles across the subsamples. We estimated summaries for protein-complexes that had four or more proteins observed in the CC or DO, after removing proteins with local pQTL (FDR < 0.5) or distal pQTL (FDR < 0.1), limiting the potential that the PC1 reflects a strong pQTL not shared by other members of the complex.

### Strain-protein outlier analysis

To identify CC strains with consistent extreme effects in both the female and male, we fit an LMM:

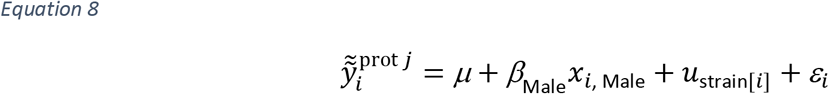

with all terms as previously defined. Effects for all CC strains for each protein 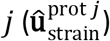 were estimated as BLUPs, which were then transformed to z-scores per protein 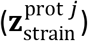. We defined a strain-protein outlier to be a protein *j* in CC strain *i* for which 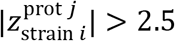. We intersected the strain-protein outliers with known CC strain-private genetic variants (Srivastava et al., 2017), identifying CC strain-private variants that likely have local effects on protein abundance. We permuted the pairings between CC strain and protein for the 6,046 strain-protein outliers to determine if the observed overlap of 69 privates was significant.

For each CC strain *i*, we defined sets of proteins that had consistently high, low, and extreme abundance based on their strain effects: 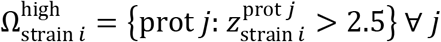, 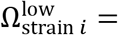 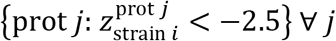, and 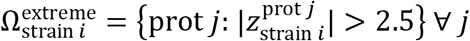, respectively. We then tested whether the CC strain-specific sets were enriched in GO and KEGG terms (*q*_set_ < 0.1), as done with proteins with sex effects.

## Data and software availability

All analyses were performed using the R statistical programming language (v3.6.1) (R Core Team, 2018). The scripts and processed data used to generate the results are available at figshare (https://doi.org/10.6084/m9.figshare.12818717). All processed data and pQTL results are available for download and interactive analysis using the QTLViewer webtool for both the CC (https://churchilllab.jax.org/qtlviewer/CC/Ferris) and DO (https://churchilllab.jax.org/qtlviewer/DO/Svenson). The mass-spec proteomics data for the CC liver samples have been deposited in ProteomeXchange (http://www.proteomexchange.org/) via the PRIDE partner repository with dataset identifier PXD018886. The mass-spec data for the DO and founder strain liver samples were previously deposited to ProteomeXchange with dataset identifier PXD002801.

## Acknowledgements

We would like to thank the members of the Churchill lab for feedback during the development of this project and in the process of composing the manuscript. We would also like to thank Lauren J. Donoghue of the University of North Carolina at Chapel Hill, John W. Keele of the United States Department of Agriculture, Paul L. Maurizio of the University of Chicago, and Bryan C. Quach of the Research Triangle Institute for their feedback on this manuscript. This work has been supported by grants from the National Institutes of Health (NIH): F32GM134599 to G.R.K.; U19AI100625, P01AI132130, and R01ES029925 to F.P.-M.V. and M.T.F.; R01GM067945 to S.P.G; and R01GM070683 to G.A.C.

## Author contributions

Conceptualization, M.T.F., S.P.G., and G.A.C.; Methodology, G.R.K., T.Z., S.P.G., and G.A.C.; Software, G.R.K, D.P., and M.V.; Investigation, G.R.K. and T.Z.; Resources, T.Z., T.A.B., P.H., G.D.S., F.P-M.V., M.T.F., and S.P.G.; Data Curation, G.R.K, T.Z., and M.V.; Writing – Original Draft, G.R.K., T.Z., and G.A.C.; Writing – Review & Editing; Visualization, G.R.K.; Supervision, S.C.M., M.T.F., S.P.G., and G.A.C.; Funding Acquisition, F.P-M.V., M.T.F., S.P.G., and G.A.C.

## Supplemental Information

Table S1. Heritability and sex effects for the proteins analyzed in the CC, DO, and founder strains, related to **Figure 1**

Table S2. Detected pQTL (LOD > 6) in the CC and DO, related to **Figures 2** and **S1**

Table S3. Consistency of haplotype effects for pQTL detected (FDR < 0.5) in the CC and DO, related to **Figures 2** and **S1**

Table S4. Mediation results for distal pQTL detected (LOD > 6) in the CC and DO, related to **Figures 2** and **S1**

Table S5. Annotated protein-complexes (Ori et al., 2016) used in this study, with added mouse ENSEMBL gene IDs and the numbers of member proteins observed in each mouse population, related to **Figures 3** and **S4**

Table S6. Summaries on heritability, pQTL, and sex effect size for individual proteins annotated to protein-complexes, related to **Figures 3** and **S4**

Table S7. Summaries of cohesiveness, heritability, and sex effect size for protein-complexes, related to **Figures 3** and **S4**

Table S8. Gene ontology enrichment results of proteins with extreme abundances specific to individual CC strains, related **Figures 7** and **S7**

Table S9. KEGG enrichment results of proteins with extreme abundances specific to individual CC strains, related **Figures 7** and **S7**

---

**Figure S1.**
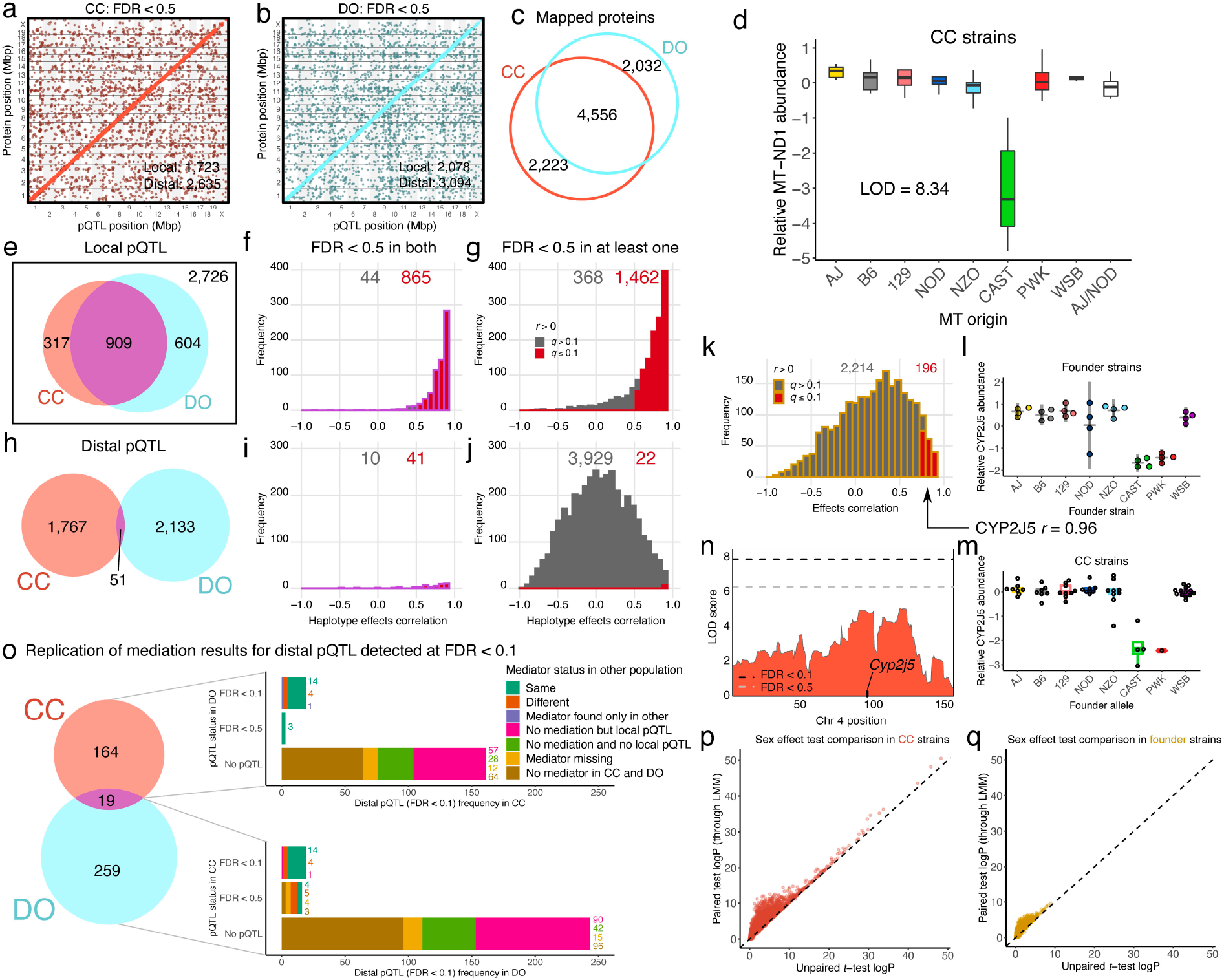
Comparison of lenient mapping results, mediation candidates, and sex effects among genetically diverse mouse populations, related to Figures 1 and 2. Leniently detected (FDR < 0.5) pQTL in the (a) CC and (b) DO. The pQTL are plotted by the genomic position of protein against their coordinate. Dot size is proportional to association strength (LOD score). (c) Venn diagram of the overlap of analyzed proteins between the CC and DO. (d) A local pQTL was detected in the CC for *mt-Nd1*, a gene encoded on the mitochondrial genome. The pQTL is characterized by low MT-ND1 in CC strains with the CAST mitochondrial genome. For six CC strains, mitochondrial inheritance was ambiguous between AJ and NOD (white boxplot). (e) Venn diagram of the overlap in local pQTL detected in the CC and DO. (f) The haplotype effects of local pQTL detected in both populations are highly consistent, as measured by the correlation coefficient comparing the effects in the CC and DO. (g) More local pQTL are consistent between the populations when also considering pQTL detected in only one of them. Red bars represent pQTL that that had significantly correlated effects (FDR < 0.1). (h) Venn diagram of overlap of distal pQTL detected in the CC and DO. (i) Using lenient detection resulted in more distal pQTL in both populations with consistent haplotype effects (41 out of 51). (j) In contrast to local genetic effects, considering distal pQTL that were leniently detected in only one of the populations resulted in many inconsistent effects comparisons, likely representing false positives or subtle distal effects specific to one population. The founder strains can reveal under-powered local pQTL in the CC (and DO), as revealed by the correlations between haplotype effects at putative local pQTL for genes with rare alleles in the CC (≤ 3 CC strains) and that did not have a local pQTL detected (FDR < 0.5) and strain effects in mice from the founder strains (k). The enrichment in positive correlations suggests the founders can provide additional information for detecting local pQTL in the CC. The gene *Cyp2j5* had significant correlation (*r* = 0.96) between the strain effects in the (l) founder strains with the (m) local haplotype effects in the CC, characterized by low CAST and PWK effects. Mean and ± 2 standard deviation bars are shown for the founder strains, and boxplots for CC strains. Four CC strains possessed the CAST allele and only one had the PWK allele, resulting in (n) poor power to detect the local pQTL of *Cyp2j5*. Stringent and lenient significance thresholds are included as horizontal black and gray dashed lines, respectively. (o) Comparison of mediation results for distal pQTL stringently detected (FDR < 0.1) in at least one of the populations. For distal pQTL detected stringently in both populations, mediation results were strongly consistent (14 out of 19). If the distal pQTL were detected leniently in the other population, mediation was still somewhat consistent (7 out of 19). When the distal pQTL was not detected in the other population, often no mediator was detected, or the candidate mediator was missing from the other population. Observance of both sexes within the (p) CC and (q) founder strains improves power to detect sex effects. The −log_10_(p-value), or logP, from a linear mixed effect model (LMM) that takes into account strain replicates compared to the logP from an unpaired t-test. The lower logP in the founder strains reflects the overall smaller sample size (32 mice compared to 116 in the CC). Dashed diagonal lines included for reference.

**Figure S2.**
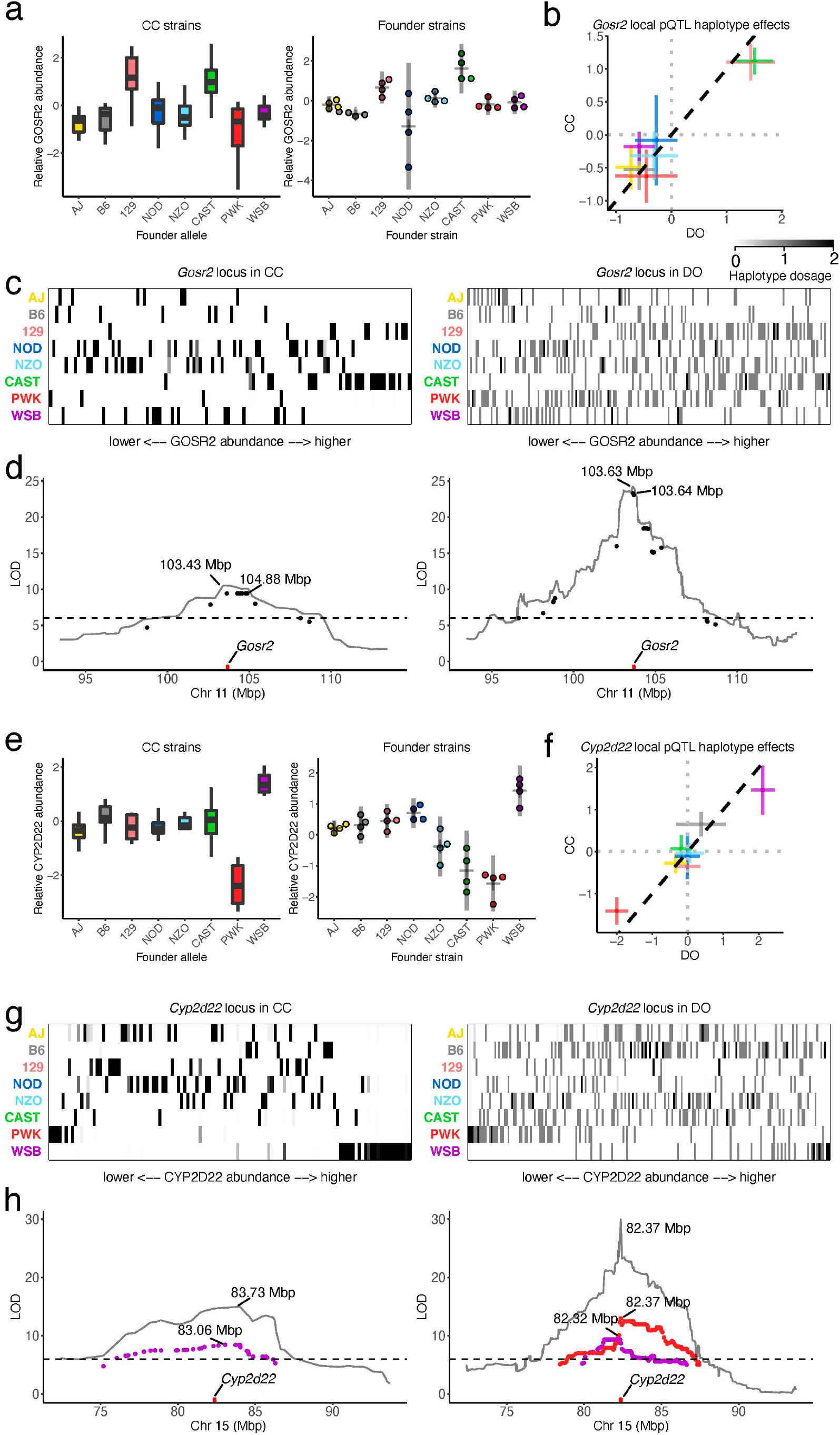
Highly consistent local pQTL effects across the CC, DO, and founder strains, related to Figure 2. Local pQTL for (a) *Gosr2* and (e) *Cyp2d22* have highly consistent effects across the CC (left) and founder strains (right). Boxplots are shown for the CC strains and mean ± 2 standard deviation bars for the founder strains. The local pQTL effects are also highly consistent between the CC and DO, both in terms of modeled effects and the actual data – for (b, c) *Gosr2* and (f, g) *Cyp2d22*. For comparisons of modeled effects (b, f), intervals represent ± standard error bars. Dashed diagonal lines included for reference. To visualize the effects in the actual data (c, g), the CC and DO data at the pQTL are represented as heatmaps with rows indicating founder allele dosage at the local pQTL (expected allele counts) and columns indicating individual mice, ordered by protein abundance. Clusters in rows towards the left or right sides represent founder haplotype effects, such as the high WSB effect in *Cyp2d22*. Haplotype-based association scans at the pQTL are overlayed with variant assocations for (d) *Gosr2* and (h) *Cyp2d22*. When the pQTL effects are approximately bi-allelic, as with *Gosr2* (d), the peak variant association and haplotype-based association are very close, consistent with a single variant largely driving the pQTL. Variants with an allele shared by only 129 and CAST, matching the effects pattern, are shown. When the effects were more complex than bi-allelic, as with *Cyp2d22* (h), there are likely multiple causal variants, and haplotype-based association produces higher scores of assocation. WSB- and PWK-private variants with LOD > 6 were included for *Cyp2d22*, highlighting LD blocks that potentially carry founder-specific variants driving their extreme effects. The larger sample size and finer mapping resolution of the DO sample are evident in the high LOD scores and narrower association peaks. Genomic positions of peak associations from variant- and haplotype-based mapping are marked. Horizontal dashed lines at LOD of 6 included as reference.

**Figure S3.**
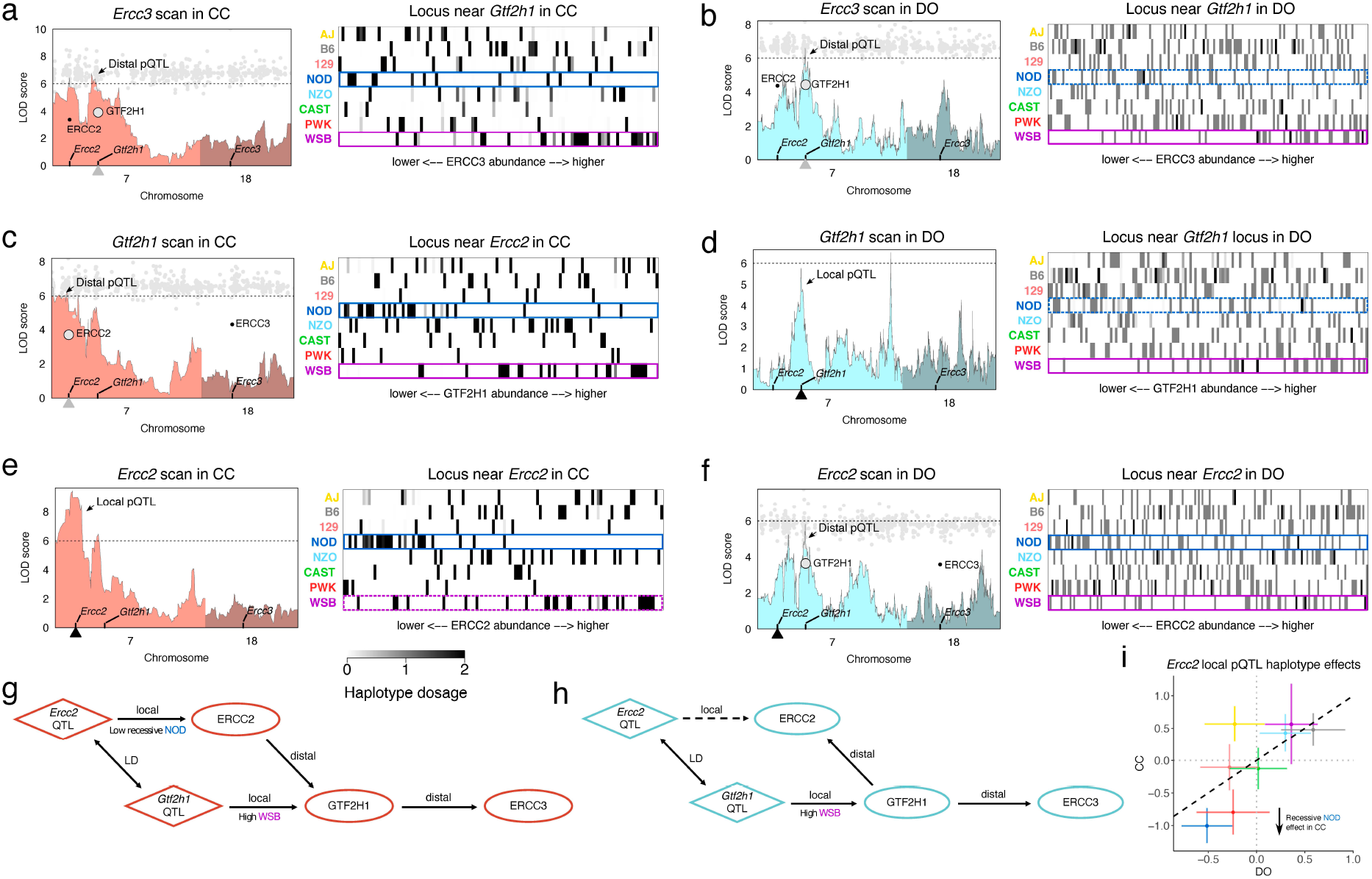
Similarity and differences in the genetic effects on *Ercc3* between the CC and DO, related to Figure 2. Suggestive distal pQTL for *Ercc3* map to chromosome 7 in both the (a) CC and (b) DO, consisting of two peaks above *Ercc2* and *Gtf2h1*, genes that are approximately 30 Mbp apart and known to interact with ERCC3. Mediation analysis identified GTF2H1 and ERCC2 as candidate drivers of the *Ercc3* pQTL. Gray dots represent mediation scores for all quantified proteins on chromosomes 7 and 18. Horizontal dashed lines at LOD of 6 included for reference. To better understand the pQTL effects, the founder haplotype inheritance was plotted as heatmaps with founder allele dosage (expected allele counts) as rows and individual mice as columns, ordered by ERCC3 abundance, at the chromosome 7 locus near *Gtf2h1*. A low NOD effect was observed, indicated by the dark blue box, which was weak to non-existent in the DO (dotted dark blue box), as well as a high WSB effect in both the CC and DO (purple boxes). (c) In the CC, *Gtf2h1* has a weak distal pQTL that mapped nearby *Ercc2* and was mediated by ERCC2 with similar low NOD and high WSB effects. (d) In the DO, *Gtf2h1* has a suggestive local pQTL (LOD < 6) and notably no suggestive association near *Ercc2*. (e) *Ercc2* has a strong local pQTL in the CC (FDR < 0.1), driven by a low NOD effect, whereas in the (f) DO, a suggestive distal pQTL is observed near *Gtf2h1*. Diagrams for the relationships defined by pQTL and mediation in the (g) CC and (h) DO. In both populations, *Ercc2* and/or *Gtf2h1* affect ERCC3 abundance, and potentially each other as well. Teasing apart the directionality of effects is further complicated by the linkage disequilibrium (LD) between *Ercc2* and *Gtf2h1*. The low NOD effect at the *Ercc2* locus is stronger in the CC, and the high WSB effect is strongest at the *Gtf2h1* locus. (i) The haplotype effects at the locus near *Ercc2* are relatively similar, though the NOD effect is more extreme in the inbred CC background. Intervals represent ± standard error bars. Dashed diagonal line included for reference.

**Figure S4.**
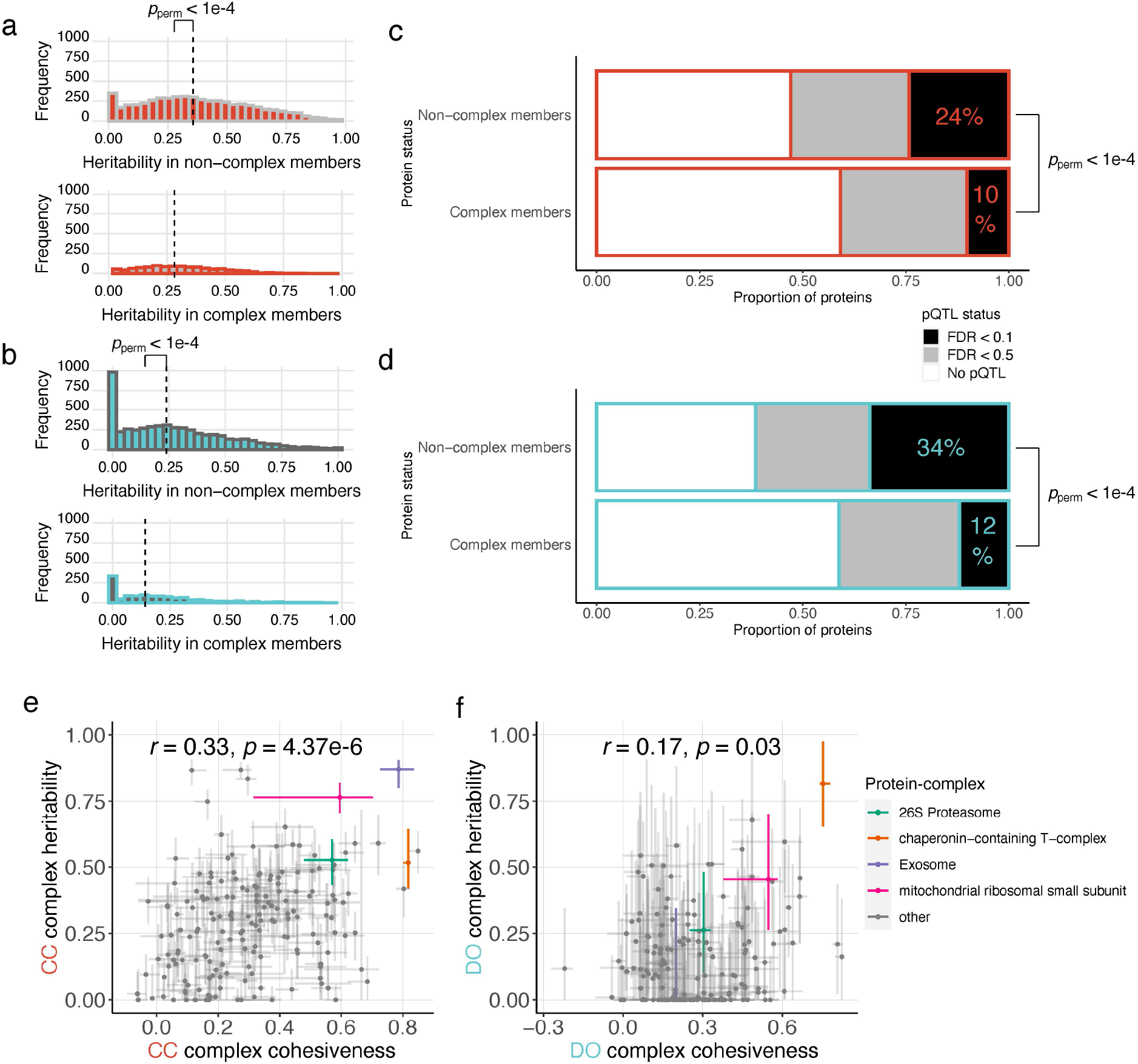
Comparatively fewer detected genetic effects on protein-complex members, and the correlation between protein-complex heritability and cohesiveness, related to Figures 3, 4, 5, and 6. Histograms of the heritability of proteins that are not annotated as members of a protein-complex (top) and that are members (bottom) in the (a) CC and (b) DO. Vertical dashed line represents the median heritability, which were significantly different (*p*_perm_ < 1e-4). A lower proportion of proteins that are members of protein-complexes possess pQTL (*p*_perm_ < 1e-4) in both the (c) CC and (d) DO. Black boxes represent stringently detected (FDR < 0.1) pQTL, gray boxes represent leniently detected (FDR < 0.5) pQTL, and white boxes represent proteins that had no pQTL detected. Numbers represent the percentage of proteins with detected pQTL at FDR < 0.1 within each category. The correlation between protein-complex heritability and cohesiveness is stronger in the (e) CC (*r* = 0.33, *p* = 4.37e-6) than the (f) DO (*r* = 0.17, *p* = 0.03). For protein-complex cohesiviness, points and bars represent medians and interquartile ranges, respectively. Protein-complex heritability was estimated from the first principal component (PC1) of the complex members, and bars represent 95% subsample intervals around the estimate. Exosome, chaperonin-containing T-complex, 26S Proteasome, and the mitochondrial ribosomal small subunit are highlighted and are examined in detail (**Figures 4, 5**, **6**, **S5**, & **S6**).

**Figure S5.**
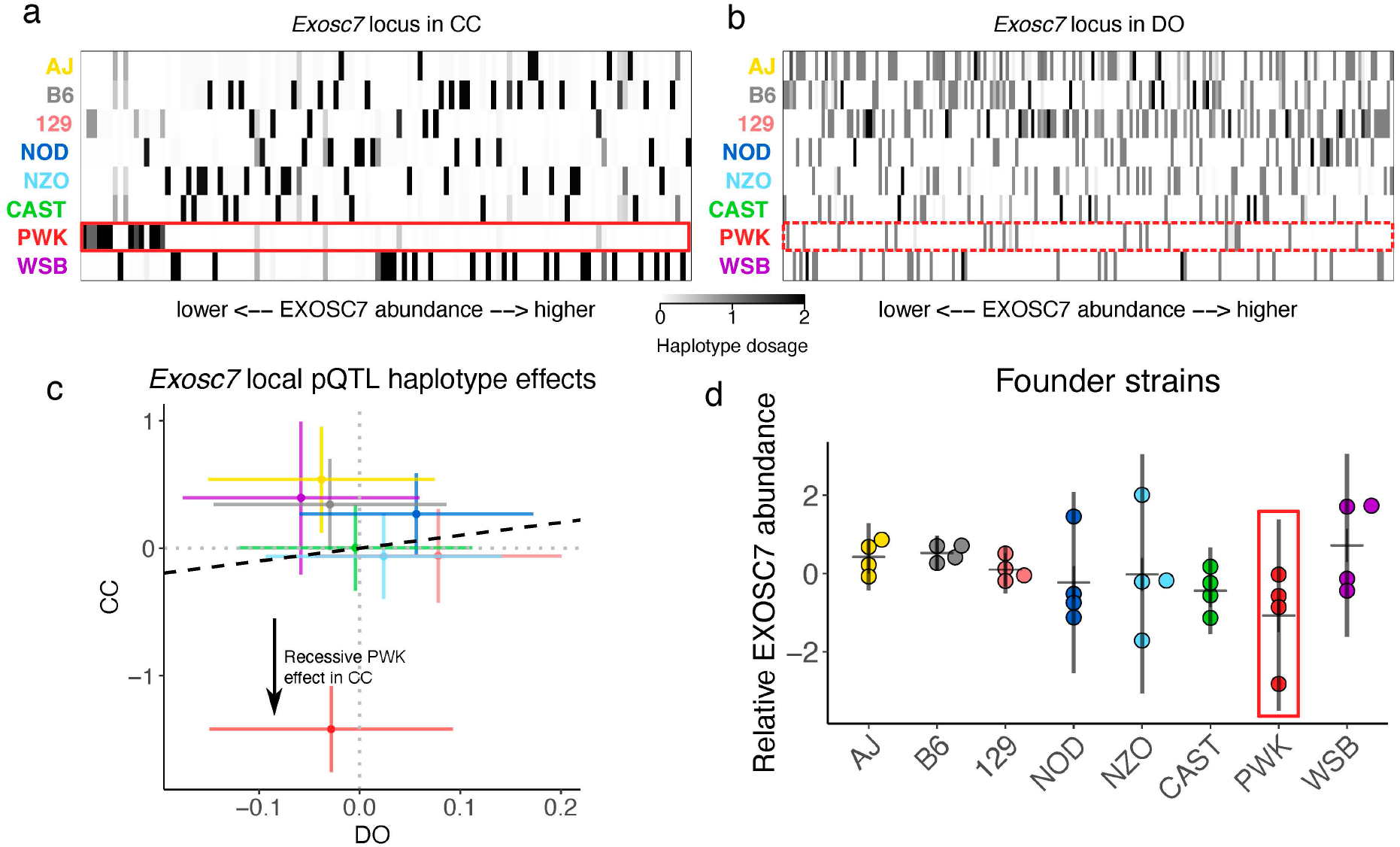
A local recessive effect on EXOSC7 that impacts the whole exosome is exposed by the inbred genomes of the CC strains, related to Figure 4. The founder haplotype inheritance at the *Exosc7* local pQTL represented as a heatmap with founder allele dosages (expected allele counts) as rows and individuals as the columns, ordered by EXOSC7 abundance, for the (a) CC and (b) DO. The cluster on the left side in the PWK row (red box) for the CC reflects the low EXOSC7 abundance observed in CC mice that are homozygous for the PWK allele at the locus. No homozygotes for PWK were observed in the DO, and the heterozygous PWK carriers do not have low EXOSC7 abundance (dotted red box). (c) Comparison of the modeled allele effects at the *Exosc7* locus reveal a marked lower PWK effect specific to the CC, which is consistent with the effects of a recessive allele. Intervals represent ± standard error bars. Dashed diagonal line included for reference. (d) Mice from the PWK strain with low EXOSC7 abundance were observed, though the effect is not as distinct as in the CC, possibly due to decreased accuracy without a bridge sample and the relative nature of the mass-spec quantification. Founder strain mice are summarized with mean ± 2 standard deviation bars.

**Figure S6.**
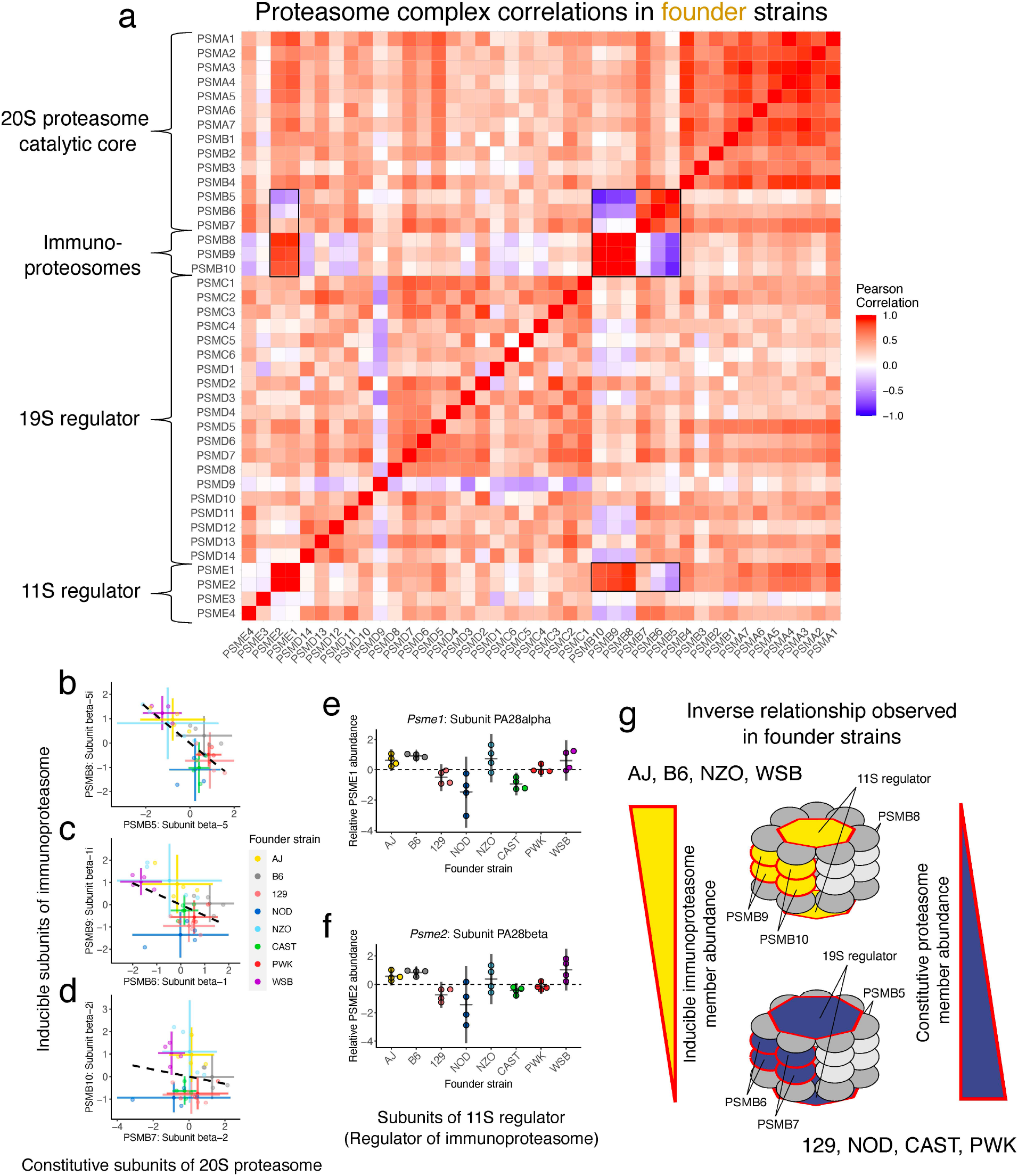
The founder strains differ in the balance between the constitutive proteasome and the inducible immunoproteasome, related to Figure 6. The correlation pattern among members of the 26S proteasome in the founder strains matches closely with those observed in the CC and DO (**Figure 6a & b**). The constitutive members of the 20S proteasome (PSMB5, PSMB6, and PSMB7) are anti-correlated with the corresponding inducible immunoproteasome components (PSMB8, PSMB9, and PSMB10) and the subunits from the 11S regulator – the regulator complex specific to the immunoproteasome – highlighted with black boxes. Founder strains with high abundance of the constitutive proteasome members have low levels of immunoproteasome proteins, and vice versa. The direction of this relationship depends on genetic factors, as revealed by the founder strains, with WSB and AJ mice consistently possessing relatively high abundance of immunoproteasome members and low abundance of the corresponding constitutive members: (b) PSMB8 vs PSMB5, (c) PSMB9 vs PSMB6, and (d) PSMB10 vs PSMB7. Intervals represent mean ± 2 standard deviation bars. Dashed lines represent the best fit line between corresponding inducible and constitutive subunits. Two of the proteins from the 11S regulator of the immunoproteasome, (e) PSME1 and (f) PSME2, have matching abundance in AJ, B6, NZO, and WSB mice as in the inducible immunoproteasome components, which is largely consistent with joint regulation of constitutive/inducible members (g). The WSB mice are consistent with genetic variation identified through QTL mapping near *Psmb9* that drives PSMB9 and indirectly PSMB6 abundance relative to each other in the recombinant CC and DO populations (**Figure 6i-l**). Founder strain mice are summarized with mean ± 2 standard deviation bars. Horizontal dashed lines provide reference at 0 for 11S regulator proteins.

**Figure S7.**
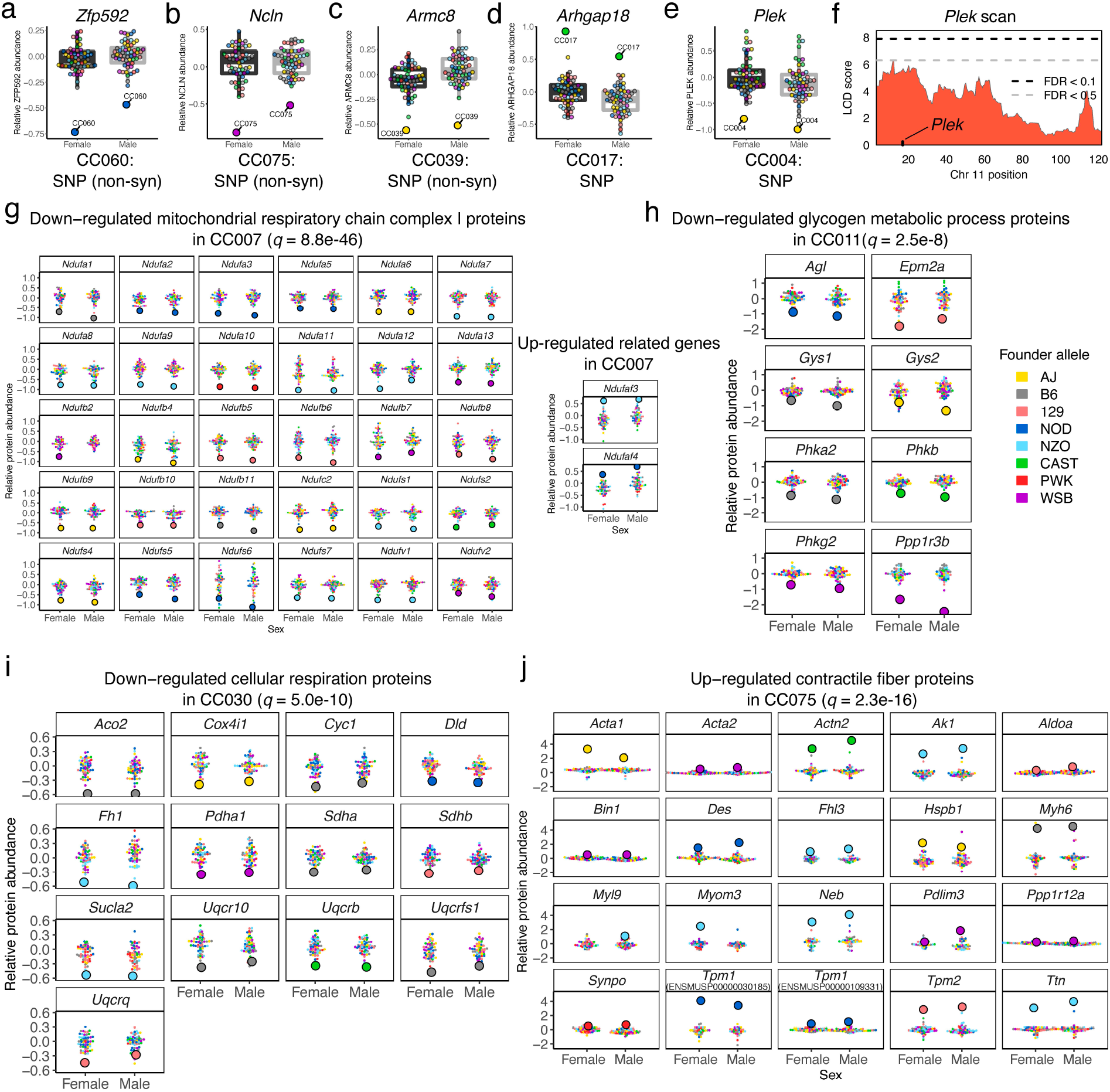
Proteins with extreme abundance in CC strains that possess unique local variants, and CC strains with extreme protein abundance patterns in functional pathways, related to Figure 7. Mutations became fixed in CC strains and were previously identified (Srivastava et al., 2017). These private variants of CC strains potentially have local effects on protein abundance. Examples include (a) *Zfp952*, (b) *Ncln*, (c) *Armc8*, (d) *Arghap18*, and (e) *Plek*. *Zfp952*, *Ncln*, and *Armc8* each possess a non-synonymous SNP variant in a CC strain which had strikingly low protein abundance. These variants may cause reduced abundance, but alternatively, they may represent a coding polymorphism in the quantified peptides, resulting in false low abundance due to the allele-specific nature of mass-spec quantification. The color or the dots indicates the founder allele at the gene. Strain-specific variants associated with high abundance at the local protein were also observed, such as high ARHGAP18 abundance in CC017. (e) A novel SNP allele in CC004 at *Plek* was associated with low abundance, potentially representing a new allele at a gene that already possessed local genetic variation from the founder strains that drove a (f) suggestive local pQTL. Stringent and lenient significance thresholds are included as horizontal black and gray dashed lines, respectively. CC strain-specific sets of proteins with extreme abundance were identified that were significantly enriched in GO and KEGG functional terms. (g) CC007 possessed distinctly low abundance of mitochondrial respiratory chain complex I proteins, as well as high abundance in two assembly factor proteins from the complex, NDUFAF3 and NDUFAF4, suggesting compensatory mechanisms may be occurring at the complex in the strain. Large dots represent the CC strain with extreme protein abundance. Other detected CC strain-specific protein dynamics include low abundance of (h) glycogen metabolic process proteins for CC0011, (i) cellular respiration proteins for CC030, and (j) high abundance of contractile fiber proteins for CC075.

## References

Albert, F.W., Bloom, J.S., Siegel, J., Day, L., and Kruglyak, L. (2018). Genetics of trans-regulatory variation in gene expression. Elife 7, 1–39.

Anderson, M.G., Smith, R.S., Hawes, N.L., Zabaleta, A., Chang, B., Wiggs, J.L., and John, S.W.M. (2002). Mutations in genes encoding melanosomal proteins cause pigmentary glaucoma in DBA/2J mice. Nat. Genet. 30, 81–85.

Ashbrook, D.G., Arends, D., Prins, P., Mulligan, M.K., Roy, S., Williams, E.G., Lutz, C.M., Valenzuela, A., Bohl, C.J., Ingels, J.F., et al. (2019). The expanded BXD family of mice: A cohort for experimental systems genetics and precision medicine. BioRxiv 672097.

Baron, R.M., and Kenny, D.A. (1986). The moderator–mediator variable distinction in social psychological research: Conceptual, strategic, and statistical considerations. J. Pers. Soc. Psychol. 51, 1173–1182.

Bates, D., Mächler, M., Bolker, B., and Walker, S. (2015). Fitting Linear Mixed-Effects Models Using lme4. J. Stat. Softw. 67.

Battle, A., Khan, Z., Wang, S.H., Mitrano, A., Ford, M.J., Pritchard, J.K., and Gilad, Y. (2015). Genomic variation. Impact of regulatory variation from RNA to protein. Science 347, 664–667.

Beasley, T.M., Erickson, S., and Allison, D.B. (2009). Rank-Based Inverse Normal Transformations are Increasingly Used, But are They Merited? Behav. Genet. 39, 580–595.

Benjamini, Y., and Hochberg, Y. (1995). Controlling the False Discovery Rate : A Practical and Powerful Approach to Multiple Testing. J. R. Stat. Soc. Ser. B 57, 289–300.

Broman, K.W., Gatti, D.M., Simecek, P., Furlotte, N.A., Prins, P., Sen, Ś., Yandell, B.S., and Churchill, G.A. (2019). R/qtl2: Software for Mapping Quantitative Trait Loci with High-Dimensional Data and Multiparent Populations. Genetics 211, 495–502.

Chesler, E.J., Lu, L., Shou, S., Qu, Y., Gu, J., Wang, J., Hsu, H.C., Mountz, J.D., Baldwin, N.E., Langston, M.A., et al. (2005). Complex trait analysis of gene expression uncovers polygenic and pleiotropic networks that modulate nervous system function. Nat. Genet. 37, 233–242.

Chick, J.M., Munger, S.C., Simecek, P., Huttlin, E.L., Choi, K., Gatti, D.M., Raghupathy, N., Svenson, K.L., Churchill, G.A., and Gygi, S.P. (2016). Defining the consequences of genetic variation on a proteome-wide scale. Nature 534, 500–505.

Churchill, G., Gatti, D., Munger, S., and Svenson, K. (2012). The Diversity outbred mouse population. Mamm. Genome 23, 713–718.

Churchill, G.A., Airey, D.C., Allayee, H., Angel, J.M., Attie, A.D., Beatty, J., Beavis, W.D., Belknap, J.K., Bennett, B., Berrettini, W., et al. (2004). The Collaborative Cross, a community resource for the genetic analysis of complex traits. Nat. Genet. 36, 1133–1137.

Collaborative Cross Consortium (2012). The Genome Architecture of the Collaborative Cross Mouse Genetic Reference Population. Genetics 190, 389–401.

Doerge, R., and Churchill, G. (1996). Permutation tests for multiple loci affecting a quantitative character. Genetics 142, 285–294.

Dudbridge, F., and Koeleman, B.P.C. (2004). Efficient Computation of Significance Levels for Multiple Associations in Large Studies of Correlated Data, Including Genomewide Association Studies. Am. J. Hum. Genet. 75, 424–435.

Elias, J.E., and Gygi, S.P. (2007). Target-decoy search strategy for increased confidence in large-scale protein identifications by mass spectrometry. Nat. Methods 4, 207–214.

Elias, J.E., and Gygi, S.P. (2010). Target-Decoy Search Strategy for Mass Spectrometry-Based Proteomics. In Methods in Molecular Biology (Clifton, N.J.), S.J. Hubbard, and A.R. Jones, eds. (Totowa, NJ: Humana Press), pp. 55–71.

French, J.E., Gatti, D.M., Morgan, D.L., Kissling, G.E., Shockley, K.R., Knudsen, G.A., Shepard, K.G., Price, H.C., King, D., Witt, K.L., et al. (2015). Diversity Outbred Mice Identify Population-Based Exposure Thresholds and Genetic Factors that Influence Benzene-Induced Genotoxicity. Environ. Health Perspect. 123, 237–245.

Gatti, D.M., Svenson, K.L., Shabalin, A., Wu, L.-Y., Valdar, W., Simecek, P., Goodwin, N., Cheng, R., Pomp, D., Palmer, A., et al. (2014). Quantitative Trait Locus Mapping Methods for Diversity Outbred Mice. G3 (Bethesda). 4, 1623–1633.

Giurgiu, M., Reinhard, J., Brauner, B., Dunger-Kaltenbach, I., Fobo, G., Frishman, G., Montrone, C., and Ruepp, A. (2019). CORUM: the comprehensive resource of mammalian protein complexes—2019. Nucleic Acids Res. 47, D559–D563.

Gralinski, L.E., Ferris, M.T., Aylor, D.L., Whitmore, A.C., Green, R., Frieman, M.B., Deming, D., Menachery, V.D., Miller, D.R., Buus, R.J., et al. (2015). Genome Wide Identification of SARS-CoV Susceptibility Loci Using the Collaborative Cross. PLoS Genet. 11, e1005504.

Gygi, J.P., Yu, Q., Navarrete-Perea, J., Rad, R., Gygi, S.P., and Paulo, J.A. (2019). Web-Based Search Tool for Visualizing Instrument Performance Using the Triple Knockout (TKO) Proteome Standard. J. Proteome Res. 18, 687–693.

Hamilton-Williams, E.E., Wong, S.B.J., Martinez, X., Rainbow, D.B., Hunter, K.M., Wicker, L.S., and Sherman, L.A. (2010). Idd9.2 and Idd9.3 Protective Alleles Function in CD4+ T-Cells and Nonlymphoid Cells to Prevent Expansion of Pathogenic Islet-Specific CD8+ T-Cells. Diabetes 59, 1478–1486.

He, B., Shi, J., Wang, X., Jiang, H., and Zhu, H. (2020). Genome-wide pQTL analysis of protein expression regulatory networks in the human liver. BMC Biol. 18, 97.

Huttlin, E.L., Jedrychowski, M.P., Elias, J.E., Goswami, T., Rad, R., Beausoleil, S.A., Villén, J., Haas, W., Sowa, M.E., and Gygi, S.P. (2010). A Tissue-Specific Atlas of Mouse Protein Phosphorylation and Expression. Cell 143, 1174–1189.

Huttlin, E.L., Bruckner, R.J., Navarrete-Perea, J., Cannon, J.R., Baltier, K., Gebreab, F., Gygi, M.P., Thornock, A., Zarraga, G., Tam, S., et al. (2020). Dual Proteome-scale Networks Reveal Cell-specific Remodeling of the Human Interactome. BioRxiv 2020.01.19.905109.

Imai, K., Keele, L., and Yamamoto, T. (2010). Identification, Inference and Sensitivity Analysis for Causal Mediation Effects. Stat. Sci. 25, 51–71.

Kang, H.M., Zaitlen, N.A., Wade, C.M., Kirby, A., Heckerman, D., Daly, M.J., and Eskin, E. (2008). Efficient control of population structure in model organism association mapping. Genetics 178, 1709–1723.

Kang, H.M., Sul, J.H., Service, S.K., Zaitlen, N.A., Kong, S.-Y., Freimer, N.B., Sabatti, C., and Eskin, E. (2010). Variance component model to account for sample structure in genome-wide association studies. Nat. Genet. 42, 348–354.

Keele, G.R., Quach, B.C., Israel, J.W., Chappell, G.A., Lewis, L., Safi, A., Simon, J.M., Cotney, P., Crawford, G.E., Valdar, W., et al. (2020). Integrative QTL analysis of gene expression and chromatin accessibility identifies multi-tissue patterns of genetic regulation. PLOS Genet. 16, e1008537.

Keller, M.P., Gatti, D.M., Schueler, K.L., Rabaglia, M.E., Stapleton, D.S., Simecek, P., Vincent, M., Allen, S., Broman, A.T., Bacher, R., et al. (2018). Genetic Drivers of Pancreatic Islet Function. Genetics 209, 335–356.

Keller, M.P., Rabaglia, M.E., Schueler, K.L., Stapleton, D.S., Gatti, D.M., Vincent, M., Mitok, K.A., Wang, Z., Ishimura, T., Simonett, S.P., et al. (2019). Gene loci associated with insulin secretion in islets from nondiabetic mice. J. Clin. Invest. 129, 4419–4432.

Kimura, H., Caturegli, P., Takahashi, M., and Suzuki, K. (2015). New Insights into the Function of the Immunoproteasome in Immune and Nonimmune Cells. J. Immunol. Res. 2015, 1–8.

Kumar, V., Kim, K., Joseph, C., Kourrich, S., Yoo, S.-H., Huang, H.C., Vitaterna, M.H., de Villena, F.P.-M., Churchill, G., Bonci, A., et al. (2013). C57BL/6N mutation in cytoplasmic FMRP interacting protein 2 regulates cocaine response. Science 342, 1508–1512.

Lippert, C., Listgarten, J., Liu, Y., Kadie, C.M., Davidson, R.I., and Heckerman, D. (2011). FaST linear mixed models for genome-wide association studies. Nat. Methods 8, 833–837.

Liu, Y., Buil, A., Collins, B.C., Gillet, L.C., Blum, L.C., Cheng, L., Vitek, O., Mouritsen, J., Lachance, G., Spector, T.D., et al. (2015). Quantitative variability of 342 plasma proteins in a human twin population. Mol. Syst. Biol. 11, 786.

Lynch, M., and Walsh, B. (1998). Genetics and Analysis of Quantitative Traits (Sunderland, MA: Sinauer Associates).

MacKinnon, D.P., Fairchild, A.J., and Fritz, M.S. (2007). Mediation Analysis. Annu. Rev. Psychol. 58, 593–614.

Marshall, R.S., and Vierstra, R.D. (2019). Dynamic Regulation of the 26S Proteasome: From Synthesis to Degradation. Front. Mol. Biosci. 6.

McAlister, G.C., Huttlin, E.L., Haas, W., Ting, L., Jedrychowski, M.P., Rogers, J.C., Kuhn, K., Pike, I., Grothe, R.A., Blethrow, J.D., et al. (2012). Increasing the Multiplexing Capacity of TMTs Using Reporter Ion Isotopologues with Isobaric Masses. Anal. Chem. 84, 7469–7478.

McMullan, R.C., Kelly, S.A., Hua, K., Buckley, B.K., Faber, J.E., Pardo‐Manuel de Villena, F., and Pomp, D. (2016). Long‐term exercise in mice has sex‐dependent benefits on body composition and metabolism during aging. Physiol. Rep. 4, e13011.

Mogil, L.S., Andaleon, A., Badalamenti, A., Dickinson, S.P., Guo, X., Rotter, J.I., Johnson, W.C., Im, H.K., Liu, Y., and Wheeler, H.E. (2018). Genetic architecture of gene expression traits across diverse populations. PLOS Genet. 14, e1007586.

Morgan, A.P., and Welsh, C.E. (2015). Informatics resources for the Collaborative Cross and related mouse populations. Mamm. Genome 26, 521–539.

Mosedale, M., Kim, Y., Brock, W.J., Roth, S.E., Wiltshire, T., Scott Eaddy, J., Keele, G.R., Corty, R.W., Xie, Y., Valdar, W., et al. (2017). Candidate Risk Factors and Mechanisms for Tolvaptan-Induced Liver Injury Are Identified Using a Collaborative Cross Approach. Toxicol. Sci. 156, kfw269.

Mosedale, M., Cai, Y., Eaddy, J.S., Corty, R.W., Nautiyal, M., Watkins, P.B., and Valdar, W. (2019). Identification of Candidate Risk Factor Genes for Human Idelalisib Toxicity Using a Collaborative Cross Approach. Toxicol. Sci. 172, 265–278.

Mulligan, M.K., Wang, X., Adler, A.L., Mozhui, K., Lu, L., and Williams, R.W. (2012). Complex Control of GABA(A) Receptor Subunit mRNA Expression: Variation, Covariation, and Genetic Regulation. PLoS One 7, e34586.

Noll, K.E., Whitmore, A.C., West, A., McCarthy, M.K., Morrison, C.R., Plante, K.S., Hampton, B.K., Kollmus, H., Pilzner, C., Leist, S.R., et al. (2020). Complex Genetic Architecture Underlies Regulation of Influenza-A-Virus-Specific Antibody Responses in the Collaborative Cross. Cell Rep. 31, 107587.

O’Brien, J.J., O’Connell, J.D., Paulo, J.A., Thakurta, S., Rose, C.M., Weekes, M.P., Huttlin, E.L., and Gygi, S.P. (2018). Compositional Proteomics: Effects of Spatial Constraints on Protein Quantification Utilizing Isobaric Tags. J. Proteome Res. 17, 590–599.

Orgel, K., Smeekens, J.M., Ye, P., Fotsch, L., Guo, R., Miller, D.R., Pardo-Manuel de Villena, F., Burks, A.W., Ferris, M.T., and Kulis, M.D. (2019). Genetic diversity between mouse strains allows identification of the CC027/GeniUnc strain as an orally reactive model of peanut allergy. J. Allergy Clin. Immunol. 143, 1027–1037.e7.

Ori, A., Iskar, M., Buczak, K., Kastritis, P., Parca, L., Andrés-Pons, A., Singer, S., Bork, P., and Beck, M. (2016). Spatiotemporal variation of mammalian protein complex stoichiometries. Genome Biol. 17, 47.

Pai, A.A., Cain, C.E., Mizrahi-Man, O., De Leon, S., Lewellen, N., Veyrieras, J.-B., Degner, J.F., Gaffney, D.J., Pickrell, J.K., Stephens, M., et al. (2012). The Contribution of RNA Decay Quantitative Trait Loci to Inter-Individual Variation in Steady-State Gene Expression Levels. PLoS Genet. 8, e1003000.

Patterson, H.D., and Thompson, R. (1971). Recovery of Inter-Block Information when Block Sizes are Unequal. Biometrika 58, 545.

Paulo, J.A., O’Connell, J.D., Everley, R.A., O’Brien, J., Gygi, M.A., and Gygi, S.P. (2016a). Quantitative mass spectrometry-based multiplexing compares the abundance of 5000 S. cerevisiae proteins across 10 carbon sources. J. Proteomics 148, 85–93.

Paulo, J.A., O’Connell, J.D., and Gygi, S.P. (2016b). A Triple Knockout (TKO) Proteomics Standard for Diagnosing Ion Interference in Isobaric Labeling Experiments. J. Am. Soc. Mass Spectrom. 27, 1620–1625.

Philip, V.M., Sokoloff, G., Ackert-Bicknell, C.L., Striz, M., Branstetter, L., Beckmann, M. a, Spence, J.S., Jackson, B.L., Galloway, L.D., Barker, P., et al. (2011). Genetic analysis in the Collaborative Cross breeding population. Genome Res. 21, 1223–1238.

Picotti, P., Clément-Ziza, M., Lam, H., Campbell, D.S., Schmidt, A., Deutsch, E.W., Röst, H., Sun, Z., Rinner, O., Reiter, L., et al. (2013). A complete mass-spectrometric map of the yeast proteome applied to quantitative trait analysis. Nature 494, 266–270.

R Core Team (2018). RSoftware2018.

Rasmussen, A.L., Okumura, A., Ferris, M.T., Green, R., Feldmann, F., Kelly, S.M., Scott, D.P., Safronetz, D., Haddock, E., LaCasse, R., et al. (2014). Host genetic diversity enables Ebola hemorrhagic fever pathogenesis and resistance. Science 346, 987–991.

Rogala, A.R., Morgan, A.P., Christensen, A.M., Gooch, T.J., Bell, T.A., Miller, D.R., Godfrey, V.L., and de Villena, F.P.-M. (2014). The Collaborative Cross as a resource for modeling human disease: CC011/Unc, a new mouse model for spontaneous colitis. Mamm. Genome 25, 95–108.

Romanov, N., Kuhn, M., Aebersold, R., Ori, A., Beck, M., and Bork, P. (2019). Disentangling Genetic and Environmental Effects on the Proteotypes of Individuals. Cell 177, 1308–1318.e10.

Shorter, J.R., Najarian, M.L., Bell, T.A., Blanchard, M., Ferris, M.T., Hock, P., Kashfeen, A., Kirchoff, K.E., Linnertz, C.L., Sigmon, J.S., et al. (2019). Whole Genome Sequencing and Progress Toward Full Inbreeding of the Mouse Collaborative Cross Population. G3 (Bethesda). 9, 1303–1311.

Sigmon, J.S., Blanchard, M.W., Baric, R.S., Bell, T.A., Brennan, J., Brockmann, G.A., Burks, A.W., Calabrese, J.M., Caron, K.M., Cheney, R.E., et al. (2020). Content and performance of the MiniMUGA genotyping array, a new tool to improve rigor and reproducibility in mouse research. BioRxiv 2020.03.12.989400.

Skelly, D.A., Czechanski, A., Byers, C., Aydin, S., Spruce, C., Olivier, C., Choi, K., Gatti, D.M., Raghupathy, N., Stanton, A., et al. (2019). Genetic variation influences pluripotent ground state stability in mouse embryonic stem cells through a hierarchy of molecular phenotypes. BioRxiv 552059.

Smith, C.M., Proulx, M.K., Lai, R., Kiritsy, M.C., Bell, T.A., Hock, P., Pardo-Manuel de Villena, F., Ferris, M.T., Baker, R.E., Behar, S.M., et al. (2019). Functionally Overlapping Variants Control Tuberculosis Susceptibility in Collaborative Cross Mice. MBio 10, 1–15.

Srivastava, A., Morgan, A.P., Najarian, M.L., Sarsani, V.K., Sigmon, J.S., Shorter, J.R., Kashfeen, A., McMullan, R.C., Williams, L.H., Giusti-Rodríguez, P., et al. (2017). Genomes of the mouse collaborative cross. Genetics 206, 537–556.

Storey, J.D., Bass, A.J., Dabney, A., and Robinson, D. (2019). qvalue: Q-value estimation for false discovery rate control.

Suhre, K., McCarthy, M.I., and Schwenk, J.M. (2020). Genetics meets proteomics: perspectives for large population-based studies. Nat. Rev. Genet.

Svenson, K.L., Gatti, D.M., Valdar, W., Welsh, C.E., Cheng, R., Chesler, E.J., Palmer, A. a, McMillan, L., and Churchill, G. a (2012). High-resolution genetic mapping using the Mouse Diversity outbred population. Genetics 190, 437–447.

Szklarczyk, D., Gable, A.L., Lyon, D., Junge, A., Wyder, S., Huerta-Cepas, J., Simonovic, M., Doncheva, N.T., Morris, J.H., Bork, P., et al. (2019). STRING v11: protein–protein association networks with increased coverage, supporting functional discovery in genome-wide experimental datasets. Nucleic Acids Res. 47, D607–D613.

Taggart, J.C., Zauber, H., Selbach, M., Li, G.-W., and McShane, E. (2020). Keeping the Proportions of Protein Complex Components in Check. Cell Syst. 10, 125–132.

Valdar, W., Holmes, C.C., Mott, R., and Flint, J. (2009). Mapping in structured populations by resample model averaging. Genetics 182, 1263–1277.

Vinayagam, A., Hu, Y., Kulkarni, M., Roesel, C., Sopko, R., Mohr, S.E., and Perrimon, N. (2013). Protein complex-based analysis framework for high-throughput data sets. Sci. Signal. 6, 1–12.

Wang, Y., Yang, F., Gritsenko, M.A., Wang, Y., Clauss, T., Liu, T., Shen, Y., Monroe, M.E., Lopez-Ferrer, D., Reno, T., et al. (2011). Reversed-phase chromatography with multiple fraction concatenation strategy for proteome profiling of human MCF10A cells. Proteomics 11, 2019–2026.

Wei, J., and Xu, S. (2016). A random-model approach to QTL mapping in multiparent advanced generation intercross (MAGIC) populations. Genetics 202, 471–486.

Williams, E.G., Wu, Y., Jha, P., Dubuis, S., Blattmann, P., Argmann, C.A., Houten, S.M., Amariuta, T., Wolski, W., Zamboni, N., et al. (2016). Systems proteomics of liver mitochondria function. Science (80-.). 352, aad0189–aad0189.

Wu, Y., Williams, E.G., Dubuis, S., Mottis, A., Jovaisaite, V., Houten, S.M., Argmann, C.A., Faridi, P., Wolski, W., Kutalik, Z., et al. (2014). Multilayered Genetic and Omics Dissection of Mitochondrial Activity in a Mouse Reference Population. Cell 158, 1415–1430.

Yang, H., Bell, T.A., Churchill, G.A., and Pardo-Manuel de Villena, F. (2007). On the subspecific origin of the laboratory mouse. Nat. Genet. 39, 1100–1107.

Yang, H., Wang, J.R., Didion, J.P., Buus, R.J., Bell, T.A., Welsh, C.E., Bonhomme, F., Yu, A.H.-T., Nachman, M.W., Pialek, J., et al. (2011). Subspecific origin and haplotype diversity in the laboratory mouse. Nat. Genet. 43, 648–655.

Yu, G., Wang, L.-G., Han, Y., and He, Q.-Y. (2012). clusterProfiler: an R Package for Comparing Biological Themes Among Gene Clusters. Omi. A J. Integr. Biol. 16, 284–287.

Zhang, J., Teh, M., Kim, J., Eva, M.M., Cayrol, R., Meade, R., Nijnik, A., Montagutelli, X., Malo, D., and Jaubert, J. (2019). A Loss-of-Function Mutation in the Integrin Alpha L (Itgal) Gene Contributes to Susceptibility to Salmonella enterica Serovar Typhimurium Infection in Collaborative Cross Strain CC042. Infect. Immun. 88, 1–19.

Zhou, X., and Stephens, M. (2012). Genome-wide efficient mixed-model analysis for association studies. Nat. Genet. 44, 821–824.

